# Tipping points are typical in ecosystems with higher-order interactions

**DOI:** 10.64898/2026.04.24.720639

**Authors:** Pablo Lechón-Alonso, Zachary R. Miller, Armun Liaghat, Paul Breiding, Mercedes Pascual, Stefano Allesina

## Abstract

Whether species-rich communities erode gradually or collapse abruptly under environmental change is a central question in ecology [1]. Classical pairwise theory predicts that coexistence is always lost gradually, through smooth declines to extinction [2], yet real ecological interactions are often strongly state-dependent – shaped by nonlinearities that fixed pairwise coefficients cannot capture [3]. Here we show that higher-order (nonlinear) interactions make abrupt, irreversible loss of coexistence a typical route to community collapse: across diverse random communities, the equilibrium supporting coexistence disappears suddenly at a fold bifurcation. Using polynomial homotopy continuation [4] to track equilibria as environmental conditions change, we find that folds progressively dominate the boundary of the coexistence domain as nonlinearity strengthens, replacing the gradual extinctions of pairwise theory. Furthermore, the sign structure of higher-order interactions controls both the onset of tipping-points and whether biodiversity buffers or amplifies collapse. Because higher-order and nonlinear interactions are intimately linked, tipping points also arise generically in pairwise models with strong nonlinearity. Applying our continuation framework to a canonical model of plant–pollinator collapse [5], we formally resolve its bifurcation structure as fold-mediated, and we show that fold bifurcations are typical across published multispecies models spanning mutualistic, competitive, and consumer-resource interactions. These results challenge the expectation that monitoring abundances suffices to anticipate collapse, and unify structural-stability theory, which delineates the safe operating space for coexistence, with critical transition theory, which characterizes the nature of its boundaries.

---

Ecosystems worldwide face environmental changes that will push communities of species beyond the conditions supporting present-day coexistence. Understanding when and how coexistence will be lost are central questions for both ecological theory and conservation [1]. Structural-stability theory addresses the first of these questions by taking a geometric approach, characterizing the size and shape of the feasibility domain – the space of environmental conditions under which a community admits an equilibrium where all species have positive abundances [2, 6–8]. In generalized Lotka–Volterra (GLV) models and related models that underpin this framework [9], interactions are pairwise and equilibrium conditions are linear in species abundances, making the feasibility domain analytically tractable. A key consequence of this linearity is that it also settles the second question: feasibility can only be lost through transcritical bifurcations, in which one or more species declines smoothly to zero abundance – a fundamentally gradual change [2].

Yet real ecological interactions are strongly state-dependent, shaped by saturating functional responses, context-dependent behaviors, environmental modification, and other nonlinearities that make per-capita interaction effects depend on community state [10–16]. Such effects cannot be absorbed into fixed pairwise coefficients [3, 17]. The theory of critical transitions shows that non-linear feedbacks can fundamentally alter how coexistence is lost, destroying stable states abruptly at fold bifurcations – tipping points – where the system shifts discontinuously and hysteretically to an alternative state [18–23]. However, this body of work has mostly relied on low-dimensional models with specific feedback architectures [24]. Although ecosystem collapse consistent with critical transitions has been explored numerically in species-rich networks with specific interaction types, such as plant-pollinator or consumer-resource models [5, 25–27], it remains unclear whether tipping points arise generically in diverse communities with nonlinear interactions.

Structural-stability theory handles diversity but assumes linearity; critical transitions theory handles non-linearity but not diversity. This gap is a key barrier to characterizing resilience in realistic communities, where it is important to understand not only the size of the safe operating space for coexistence, but also how collapse unfolds once its boundary is crossed [24, 28, 29].

Here we address this question by extending the GLV model with quadratic higher-order interaction (HOI) terms as a minimal representation of state-dependent interaction effects [17]. The resulting equilibrium conditions are systems of coupled multivariate polynomials of degree two, for which no closed-form solution exists in general beyond small systems [30]. In addition, the exponential growth in the number of equilibria makes feasibility boundaries challenging to study even computationally [31]. We overcome these obstacles using polynomial homotopy continuation[4], a method from numerical algebraic geometry [32] that tracks all solutions of polynomial systems as parameters vary.

By subjecting model communities with HOIs to growth-rate perturbations and mapping their feasibility boundaries, we show that fold bifurcations progressively dominate the boundary as nonlinearity strengthens, replacing the gradual extinctions that characterize pair-wise theory. The sign structure of HOIs controls tipping-point prevalence and whether biodiversity acts as a buffer or amplifier of abrupt collapse. Furthermore, because saturating functional responses generate effective higher-order interactions [33, 34], we use the same continuation framework to explore tipping points in pairwise models with nonlinear interactions. We formally resolve the bifurcation structure of a canonical model of ecosystem collapse in plant-pollinator networks, confirming it is fold-mediated. More broadly, we find that fold-mediated coexistence loss is common in other published multispecies models across interaction types and functional forms, and becomes more common as interactions are more nonlinear.

## Two routes to coexistence loss

We study coexistence under environmental perturbations in generalized Lotka-Volterra (GLV) models with higher-order interactions (HOIs),

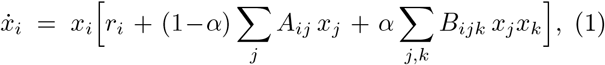

where **x** = (*x*_*i*_) are abundances, **r** = (*r*_*i*_) are intrinsic growth rates, *A*_*ij*_ entries are pairwise interactions, *B*_*ijk*_ entries are three-way interactions (HOIs), and *α* ∈ [0, 1] controls the relative weight of higher-order vs. pairwise terms (recovering the classic pairwise GLV when *α* = 0; see [35]). Throughout, we model environmental change by perturbing intrinsic growth rates along fixed directions:

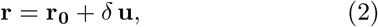

where **u** is the direction of the perturbation and *δ* is its magnitude. We explore the robustness of the coexistence equilibrium and how it is lost by tracking the long-term community state as *δ* varies (see SI Sec. 2). In the pairwise model (*α* = 0), the coexistence equilibrium **x**^*^ is mapped linearly to **r**, so it shifts continuously under environmental forcing, and extinctions can only occur when species abundances are driven smoothly to zero. With HOIs, however, the same ecosystem can respond in fundamentally different ways depending on the perturbation direction, even from the same baseline coexistence state (Fig. 1, *δ* = 0, and SI Sec. 1). Along one direction (Fig. 1**a**), numerical integration of Eq. (1) (crosses) tracks the stable coexistence branch found by polynomial homotopy continuation (lines) until one species smoothly declines to zero abundance at a transcritical bifurcation (*δ*_*c*_, vertical dashed line); reversing the perturbation recovers coexistence at the same threshold. Along a different direction (Fig. 1**b**), we find instead a critical transition: trajectories follow the stable coexistence branch until it terminates at a fold (saddle-node) bifurcation, beyond which no nearby stable coexistence equilibrium exists and the system collapses to an alternative attractor where one species is absent. This transition exhibits hysteresis: Co-existence is regained only after reversing the perturbation well past the forward critical point (Fig. 1**b**).

**FIG. 1.**
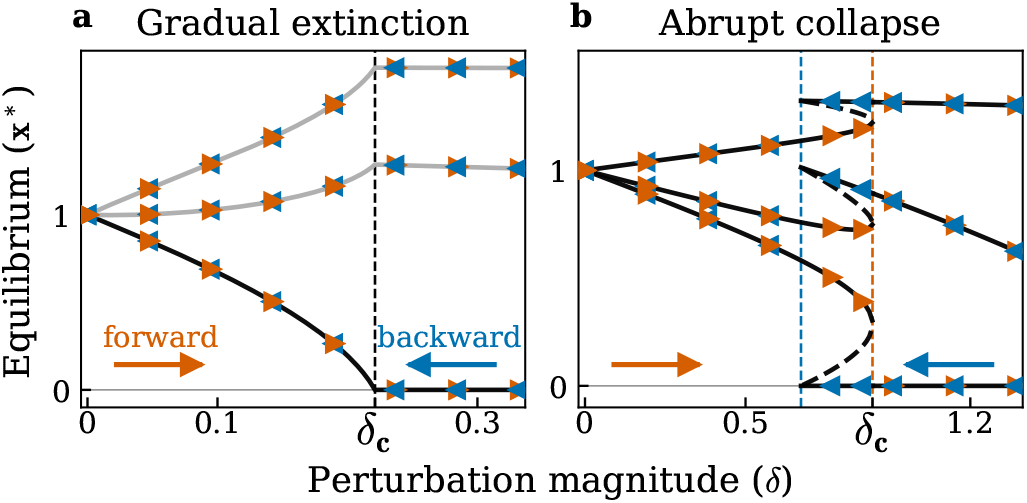
Two ways coexistence can be lost in a three-species community with higher-order interactions. Black/grey curves show equilibrium abundances of each species computed using polynomial homotopy continuation (**x**^⋆^, y-axis) as perturbation magnitude (*δ*, x-axis) varies along two fixed perturbation directions (panels). Line type indicates stability of the equilibrium (solid: stable; dashed: unstable). Triangles mark endpoints of numerical integrations of Eq. (1) that validate the continuation curves: in the forward sweep (orange right-pointing triangles), *δ* is increased in small steps from *δ* = 0, with each integration initialized at the final state of the previous step; the backward sweep (blue left-pointing triangles) reverses the process from the post-collapse state, testing where coexistence is recovered. Vertical dashed lines mark critical perturbations where coexistence is lost/gained. Panel **a** shows gradual extinction: one species declines continuously to zero abundance at *δ*_c_ (where a transcritical bifurcation takes place), and the same threshold is recovered upon reversal of the perturbation (arrows). Panel **b** shows abrupt collapse: coexistence is lost at a fold bifurcation as the perturbation magnitude increases. When the perturbation is gradually reversed (backward sweep), recovery occurs at a different critical value, indicating hysteresis.

## Higher-order effects reshape coexistence

To connect the bifurcation picture in Fig. 1, which follows two specific perturbation directions, to a geometric, global notion of robustness, we next visualize coexistence geometry in a two-species system. With two species, interactions are necessarily pairwise, but they include non-linear per capita terms. Figure 2 shows how the feasibility domain – the set of growth rates that admit a positive coexistence equilibrium – deforms as non-linearity (*α*) strengthens (panels **a**-**e**). For *α* = 0 the domain is the classical feasibility cone [6–8] whose boundaries are transcritical bifurcations – gradual extinctions at which one species approaches zero continuously (Fig. 1**a**). As *α* increases, these gradual boundaries bend (Fig. 2**b**), and for sufficiently strong HOIs, a new boundary type emerges: the coexistence equilibrium abruptly disappears through the loss of real solutions to the equilibrium equations at a fold bifurcation (Fig. 2**c**-**e**). This abrupt boundary initially lies outside the feasibility domain for small *α* but ultimately truncates it. Throughout, we restrict our attention to the focal coexistence branch, which continues smoothly from the unique coexistence equilibrium at *α* = 0. Other equilibria can be created by HOIs, generating additional interior boundaries (e.g., interior black curves in panels **c**-**e**).

**FIG. 2.**
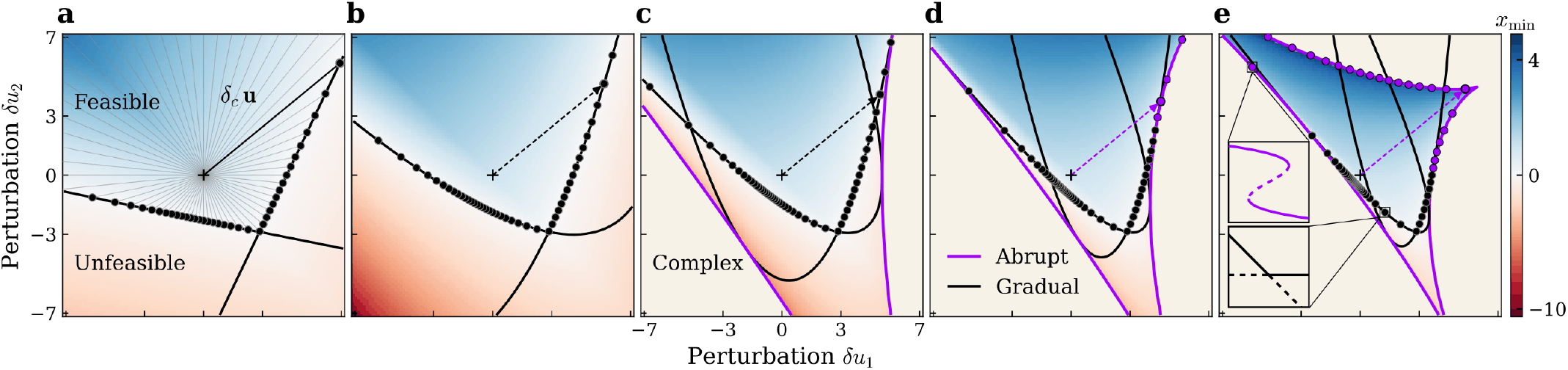
Feasibility domains as HOI strength increases. Shading shows min_*i*_ 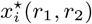 along the focal coexistence equilibrium branch 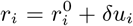. Blue indicates a positive equilibrium (feasible), red indicates at least one negative component (unfeasible), off-white background indicates regions where no real equilibrium exists (complex). Black curves are boundaries where a species gradually declines to zero at a transcritical bifurcation (gradual boundary, see bottom inset in panel **e**, and Fig 1**a**). Purple curves are boundaries where two equilibria collide and disappear from the real line at a fold bifurcation (abrupt boundary, see top inset of panel **e**, and Fig 1**b**). Left to right, stronger HOIs bend gradual boundaries (**a, b**) and generate abrupt boundaries (**c**-**e**) that ultimately truncate the feasible region. The baseline environmental state (cross) is plotted across all panels, and one perturbation direction **u** is highlighted (dashed segment). Along this direction, the distance to the gradual boundary (*δ*_c_) varies as *α* increases (**a**-**c**), and eventually the gradual boundary is superseded by an abrupt one (**d**,**e**). Lines: analytic boundaries obtained using elimination theory; points: numerical boundary points obtained by polynomial homotopy continuation along a set of 100 perturbation directions. Interior black curves correspond to gradual boundaries of additional equilibria introduced by HOIs, which do not affect the focal branch.

In two dimensions, this geometric picture can be computed in closed form: gradual boundaries follow from the vanishing of a coordinate of the coexistence equilibrium, while abrupt boundaries correspond to the vanishing of the appropriate discriminant (a classical algebraic object called tact invariant [36]; see Methods and SI Sec. 3). However, the resulting expressions are already un-wieldy for *n* = 2 and become intractable in higher dimensions, motivating our computational approach. We use the analytic boundary in Fig. 2 as a ground-truth benchmark to validate the continuation-based numerical pipeline used for larger communities: the numerical boundary points found using polynomial homotopy continuation along fixed perturbation directions (dots, see Eq. (2)) coincide with the exact analytic curves (lines).

We next apply this numerical boundary-tracking algorithm to HOI communities with *n >* 2 species (Fig. 3), systematically varying HOI strength *α*, community size *n*, and the mean of *B*_*ijk*_ (*µ*_*B*_) to probe competitive, mixed, and facilitative HOI regimes (Fig. 3**g**). To isolate the effect of nonlinearity on robustness, we construct random communities that share the same locally stable baseline equilibrium across all *α* (Methods), perturb growth rates along many directions, and classify the first feasibility boundary encountered as gradual or abrupt. Loss of stability before feasibility occurs very rarely (SI Appendix E) and is omitted for clarity.

**FIG. 3.**
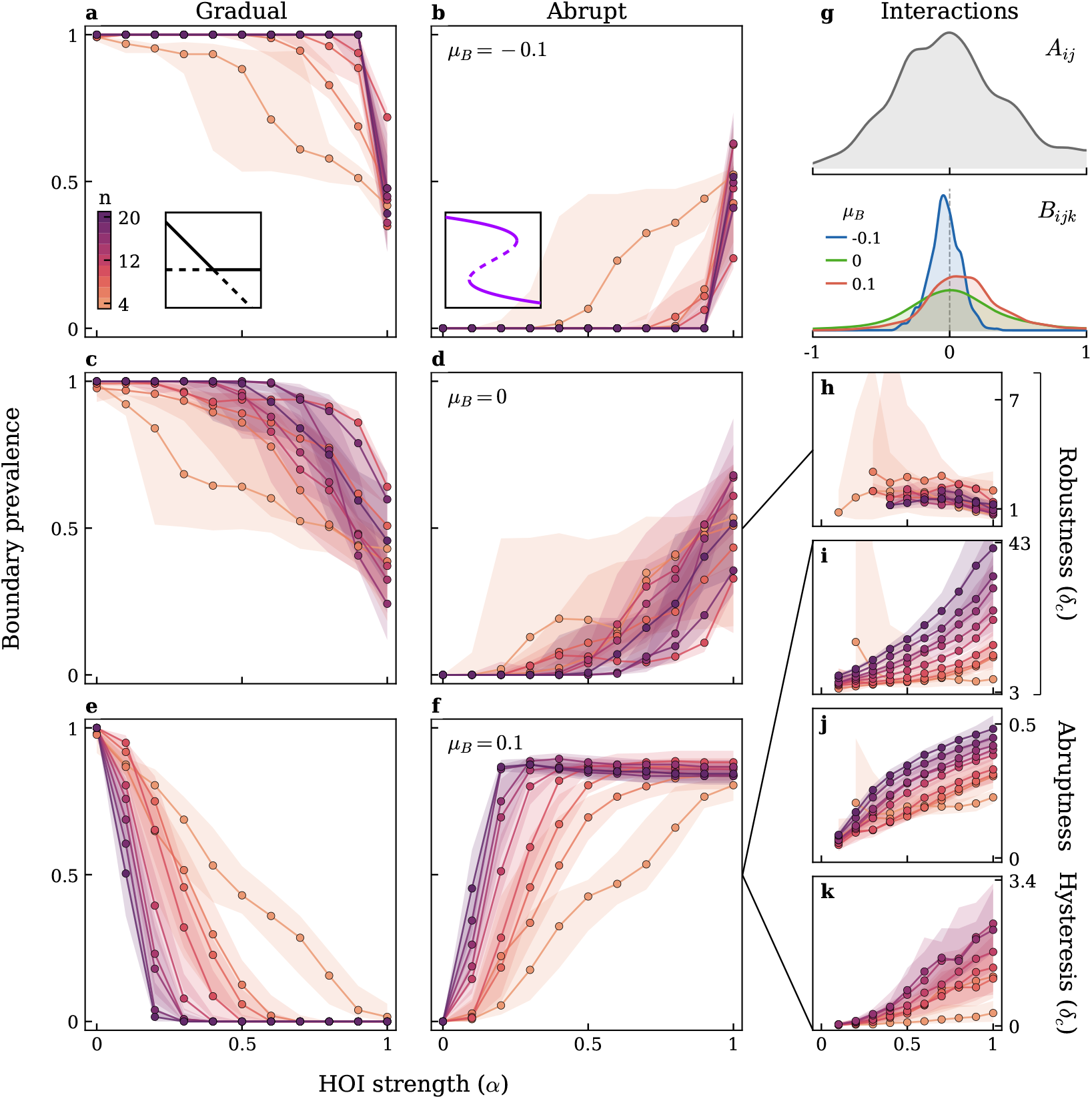
The strength and sign of higher-order interactions controls tipping-point prevalence. Panels show results for random multispecies communities (*µ*_*A*_ = 0, *σ*_*A*_ = *σ*_*B*_ = 1) with three levels of HOI mean (*µ*_*B*_ ∈ {−0.1, 0, 0.1}, rows). Panel **g** shows the empirical distributions of off-diagonal pairwise interactions *A*_*ij*_ (shared across *µ*_*B*_, grey) and higher-order interactions *B*_*ijk*_ (coloured by *µ*_*B*_) for communities of size *n* = 4. For each HOI strength (*α*, x-axis) and community size (*n*, colors), we perturb growth rates along 128 directions and classify the first boundary encountered by the continued coexistence equilibrium. Panels **a**-**f** show the fraction of directions terminating at a gradual (left column) or abrupt (right column) boundary, for each value of *µ*_*B*_ (row). Connected points give the median across replicates for each (*n, α*) condition. Panels **h**-**k** characterize the abrupt boundaries: (**h, i**) critical distance to the boundary *δ*_c_ for *µ*_*B*_ = 0 (**h**) and *µ*_B_ = 0.1 (**i**); (**j**) minimum species abundance at the equilibrium just before crossing the boundary 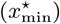 (**k**) amount of *δ* that must be reversed before ecosystem recovery after collapse–zero indicates immediate recovery, values *>* 0 indicate hysteresis. (**j**) and (**k**) show results for *µ*_*B*_ = 0.1. Note that in contrast to the competitive and mixed regimes (rows 1-2), increasing diversity in the facilitative regime (row 3) amplifies fold prevalence and community robustness as a function of *α*. Cases where no boundary was reached or where stability is lost before feasibility are infrequent for all *α* and *n*, and are omitted for clarity.

For all combinations of *n* and *µ*_*B*_, abrupt boundaries become more likely as *α* increases. When HOI coefficients are drawn symmetrically around zero (*µ*_*B*_ = 0; Fig. 3**c**,**d**), fold boundaries progressively replace transcritical extinctions as the typical mode of coexistence failure at moderate to high *α*. This basic pattern is robust to alternative parameterizations that relax our imposed controls, including the planted equilibrium, convex interpolation between pairwise and HOI terms, and the imposition of baseline stability through strong self-regulation (see SI Sec. 6).

The shift from gradual to abrupt boundaries is greatest when HOI coefficients are biased toward positive (facilitative) values (*µ*_*B*_ = 0.1; Fig. 3**e**,**f**): fold boundaries dominate even at moderate *α*, and the transition to the abrupt regime occurs earlier and more completely than in the mixed case. Strikingly, the effect of biodiversity is reversed in this regime. Whereas increasing community size *n* reduces fold prevalence in the competitive and neutral regimes – diversity buffers against abrupt extinctions – facilitative HOIs cause richer communities to be more prone to tipping-point collapse (Fig. 3**f**). These three qualitative patterns – the monotonic rise of fold prevalence with *α*, the earlier onset under facilitative HOIs, and the sign-reversal of the diversity effect across *µ*_*B*_ – persist when the mean pairwise interaction *µ*_*A*_ is also varied across competitive, neutral, and facilitative values (SI Sec. 4, Fig. S3).

Panels **h**-**k** further characterize the abrupt boundaries in the *µ*_*B*_ ≥ 0 regime (results for all regimes in SI Appendix E). In the facilitative regime (*µ*_*B*_ = 0.1) richer communities encounter more tipping point boundaries yet are farther from them in parameter space (panel **i**) – a tradeoff between likelihood and imminence that is absent in the mixed regime (*µ*_*B*_ = 0; panel **h**), where *δ*_*c*_ is only weakly related to *α* for all community sizes. With increasing *α*, the species that goes extinct at the boundary retains higher abundance immediately before the tipping point is reached (panel **j**), indicating a more abrupt transition. Additionally, hysteresis strengthens (panel **k**): the perturbation must be reversed further before coexistence recovers.

When HOI coefficients are instead biased toward negative (competitive) values (*µ*_*B*_ = −0.1; Fig. 3**a**,**b**), fold boundaries are largely suppressed: while abrupt extinctions occur, especially in small communities, coexistence is primarily lost through gradual extinctions even at high *α*. The suppression of folds in this regime raises a natural question: Are there conditions under which fold boundaries are ruled out entirely? A fold bifurcation requires two equilibria to collide and annihilate; if the coexistence equilibrium is globally asymptotically stable on the positive orthant, it is the only interior equilibrium and such collisions are precluded. Using a Lyapunov function [37, 38], we derive sufficient conditions on *A* and *B* that guarantee global stability and thereby rule out fold boundaries (Methods, and SI Sec. 5). This result extends to the HOI setting a problem studied by Cenci and Saavedra [9] for nonlinear pairwise models: when do the equilibrium conditions remain topologically equivalent to the linear case, preserving uniqueness of the coexistence equilibrium (and precluding tipping points)?

## Abrupt boundaries in pairwise models with nonlinearities

We used a random GLV+HOI model to explore whether fold bifurcations become typical when interactions are strongly state-dependent. Effective HOIs arise in any model with nonlinear interactions [33, 34, 39], with a non-random structure that reflects that underlying interaction mechanism(s). To test whether fold-mediated collapses also occur in such models, we applied our continuation framework to a canonical model of ecosystem collapse in plant-pollinator networks [5], which features mutualistic interactions mediated by saturating functional responses. These saturating responses generate higher-order effective interactions (SI Sec. 6), providing a concrete, mechanistic instance of the polynomial nonlinearity found in our GLV+HOI model. We implement the model for 10 plants and 10 pollinators following the original parameterization in [5] and apply our perturbation protocol. Using polynomial homotopy continuation, we map out the bifurcation where the system collapses (Fig. 4**a**; see SI Sec. 6 for details), confirming that it encounters a fold bifurcation, with hysteresis delimited by the endpoints of the stable and unstable branches, as in Fig. 1**b**. This example illustrates the power of our homotopy continuation approach to resolve the structure of bifurcations in complex ecological models: the collapse in this model has long been conjectured to be a tipping point, but establishing this formally is challenging using only simulations of the model dynamics [28].

**FIG. 4.**
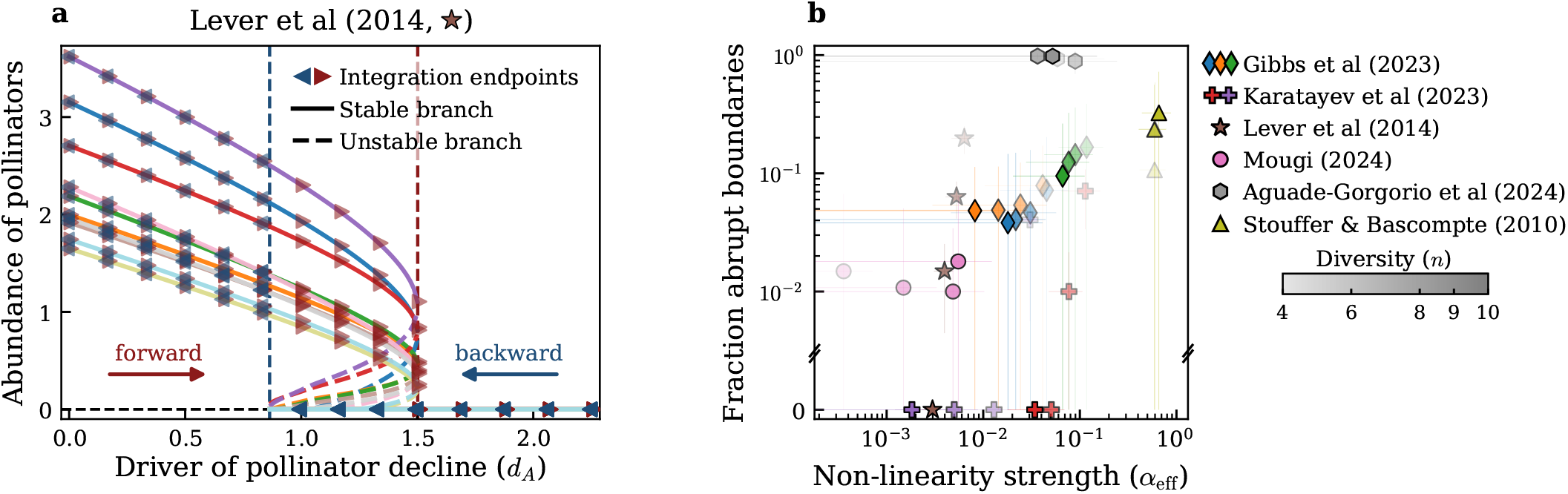
Fold-mediated collapse is common in published multispecies ecological models. **(a)** Bifurcation diagram for the canonical plant-pollinator model of Lever et al. (2014). Polynomial homotopy continuation tracks two equilibrium branches (stable, solid; unstable, dashed) as pollinator mortality *d*_*A*_ increases. The forward sweep (dark red triangles) shows numerical integration endpoints confirming that the realized community state follows the stable branch until a fold bifurcation, where stable and unstable branches meet and annihilate (dark red dashed vertical line). Beyond the fold, the system collapses to a zero-abundance state. The backward sweep (dark blue triangles) shows numerical integration from the collapsed state as the perturbation is gradually reduced, revealing hysteresis. **(b)** Fraction of perturbation directions terminating at a fold (abrupt) boundary versus effective nonlinearity strength *α*_*eff*_ across published multispecies ecological models (shapes). Shading indicates species richness. Diversity effects on fold prevalence are model-dependent: fold prevalence increases with richness in some systems (Aguadé-Gorgorió et al., Stouffer & Bascompte) but decreases in others (Lever et al.). Error bars show interquartile ranges across replicate parameterizations. Despite spanning different interaction types, functional forms, and parameterization schemes, models with stronger effective nonlinearity tend to exhibit a higher prevalence of fold-mediated coexistence loss, matching the pattern from our random HOI ensembles (Fig. 3).

To test whether tipping points are typical across other nonlinear interaction models, we repeated our perturbation protocol on a range of published multispecies models spanning competitive HOIs [40], consumer-resource systems [27], food webs with ecosystem-engineering effects [41], mixed competitive-facilitative communities [42], and empirical trophic networks [43] (Fig. 4**b**, and SI Sec. 6). Despite the diverse origins and interaction types in these models, the results are consistent with our random ensembles: models with stronger effective nonlinearity (*α*_*eff*_, SI Sec. 6) exhibit a high fraction of fold-mediated boundaries. In line with our finding that diversity can increase or decrease the prevalence of fold bifurcations depending on the sign of HOIs, the effect of diversity varies across models.

## Discussion

Our results show that higher-order interactions make fold-mediated, tipping-point loss of coexistence a common route to community collapse. We find the same outcome using alternative HOI sampling approaches [40] and in other nonlinear models (SI Sec. 6). Fig. 4**b** shows that fold-mediated collapse arises in multispecies models spanning mutualistic plant-pollinator networks [5, 43], consumer-resource systems with nonlinear functional responses [27], and communities with adaptive foraging [41] or multiple interaction types [42]. This cross-model comparison suggests that tipping points are a widespread consequence of nonlinear state-dependent interactions in complex communities.

We further found that the mean HOI effect controls important features of the transition from gradual to abrupt extinctions. HOIs that are facilitative on average make fold boundaries dominant even at weak nonlinearity, whereas competitive HOIs largely suppress them. Tipping points in ecological models have often been associated with positive feedback loops, such as in mutualistic systems [19, 20]. The facilitative HOI regime mirrors this type of mechanism, because HOIs tend to weaken antagonistic pairwise effects (and amplify mutualistic ones). HOIs contribute more strongly to net growth rates when most species are relatively abundant, while pairwise effects should dominate when abundances are lower. This creates the conditions for multistability, driven by a diffuse positive feedback loop between abundances and growth rates – HOIs can rescue the community from extinctions driven by pairwise interactions, but only once abundances are sufficiently high. Consistent with this interpretation, we found that HOIs simultaneously increase the prevalence of tipping points and the distance to a coexistence boundary (Fig. 3**i**), postponing extinctions compared to the pairwise baseline (consistent with experimental evidence [44]). Multistability, in turn, creates the conditions for a fold bifurcation (stable and unstable equilibria colliding and annihilating). However, this difference in tipping-point prevalence is driven by subtle changes in the mean HOI effect (Fig. 3**g**). Even in the facilitative regime, many individual HOIs are negative, suggesting that specific interaction structures are not necessary for tipping points to become common. Furthermore, restricting interactions to be only negative (purely competitive, SI Sec. 4) does not change the increased prevalence of folds with α.

Our analysis focused throughout on the focal equilibrium that is continuous with the unique coexistence equilibrium in the pairwise GLV model, but HOIs generically create additional coexistence states [31, 45, 46]. This implies that a complete picture of community robustness must account for both the size and shape of the local feasibility domain around each equilibrium, but also the total number and distribution of equilibria. Accounting for this global picture, nonlinearities might make any particular community more prone to tipping abruptly, but increase the chances that the community shifts to another configuration with high richness.

We also focused on loss of equilibrium feasibility; however, coexistence can alternatively be lost via instability, for example mediated by delayed feedbacks [47] or explicit time delays [48]. Characterizing when each route dominates is an important open question [49].

Our analysis leveraged polynomial homotopy continuation to navigate the nonlinear feasibility domains that arise in large ecological communities with nonlinear and nonpairwise interactions. This numerical technique has significant potential to extend the structural-stability perspective beyond pairwise models and help unify our understanding of the regime shifts that occur in complex ecological models. We may also gain a deeper understanding of these systems by looking across fields: ‘explosive’ discontinuous transitions have been reported in higher-order network models of contagion [50, 51], synchronization [52], and social dynamics [53], typically derived from mean-field reductions. Whether these share a formal connection with the species-resolved fold bifurcations we identify is an intriguing parallel that warrants further investigation.

Taken together, our findings show that HOIs add more than quantitative corrections to pairwise coexistence theory. They qualitatively transform the geometry of co-existence, replacing gradual erosion of coexistence with abrupt hysteretic collapse as the generic mode of community failure. This perspective provides a missing link between structural-stability theory, which quantifies how large the safe operating space is, and critical-transitions theory, which determines what happens when its boundary is crossed. By mapping how the strength and sign of nonlinear interactions govern tipping-point prevalence across diverse community types, our results establish a baseline expectation for when abrupt ecosystem collapse may become typical.

## MATERIALS AND METHODS

### Polynomial Homotopy Continuation

Following Eq. (1) we define the coexistence equilibrium as any real solution 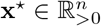 of

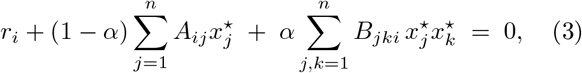

for 1 ≤ *i* ≤ *n*. When we perturb the growth-rate as **r**_0_ + *δ***u** (see Eq. (2)) we obtain from (3) the following system of polynomial equations depending on both **x** and *δ*:

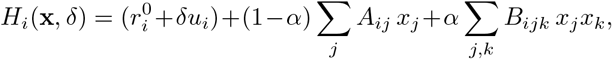

The coexistence equilibrium **x**^⋆^ corresponding to *H*(**x**, *δ*) = 0 depends continuously on *δ*. Therefore, if we move *δ* in a continuous way, **x**^⋆^ will move along. This observation is the basis of polynomial homotopy continuation [32].

If 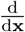 *H* is invertible at (**x**, *δ*), the implicit function theorem implies that **x** = **x**(*δ*) is locally a differentiable function of *δ*. Implicit differentiation yields

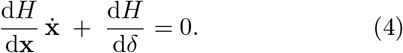

We always have the initial equilibrium **x**^⋆^ = **1** corresponding to *δ* = 0, by construction (see Parameter sampling protocol, below). Together with Eq. (4) this defines an initial value problem, which can be solved using numerical integration. All of this is implemented to be used out-of-the-box in the software HomotopyContinuation.jl [4], which we use in our work.

### Parameter sampling protocol

To isolate the effect of HOI strength α on the geometry of feasibility boundaries, we construct random communities that share a common baseline coexistence state across all *α*. Off-diagonal pairwise entries are drawn as 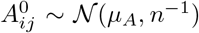 and HOI entries as 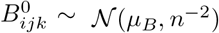, where the variance scalings ensure that both pairwise row sums and HOI slice sums contribute *O*(1) to per-capita growth rates as community size grows. Diagonal entries are then set so that row sums of A and slice sums of *B* each equal −1, planting a coexistence equilibrium **x**^⋆^ = **1** corresponding to **r** = **1** for every *α* ∈ [0, 1]. Local stability is guaranteed by rescaling each matrix by its symmetrised spectral radius and subtracting a unit diagonal, so that the symmetrised pairwise and HOI sectors (*S*_*A*_ and 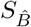) are each negative definite. Because the Jacobian at **x**^⋆^ is a convex combination of these two sectors, Hurwitz stability holds for all *α*. For each community realisation, the same bank of perturbation directions {**u**_*k*_}, sampled uniformly on the unit sphere, is used across all *α*, so that changes in boundary-type frequency reflect reshaping of the feasibility domain rather than a changing reference state or perturbation set. See SI Sec. 4 for details.

### Boundary detection with homotopy continuation

For each perturbation direction **u**, we track the coexistence equilibrium from *δ* = 0 to *δ*_max_ using coefficient-parameter homotopy continuation implemented in `HomotopyContinuation.jl` [4]. At each continuation step we monitor the tracked solution for three boundary events: (i) a gradual boundary, where the minimum species abundance drops below *x*_tol_ = 10^−12^, signalling a transcritical bifurcation; (ii) an abrupt boundary, where the tracker terminates before any component reaches *x*_*tol*_, signalling a fold bifurcation at which the Jacobian 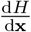 becomes singular and the real equilibrium branch ceases to exist; and (iii) an unstable boundary, where the leading eigenvalue of the community Jacobian *J* = *D*(**x**^⋆^) *J*_*F*_ crosses *λ*_tol_ = 10^−9^ while all abundances remain above *x*_*tol*_. Along each direction, we record the type of the boundary and the value *δ*_*c*_. Complete details on the algorithmic approach are given in SI Sec. 2.

### Computing the feasibility boundaries analytically for

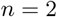

To compute the gradual and abrupt boundary analytically, we substitute in (3) 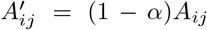 and 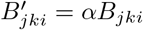, and obtain polynomials

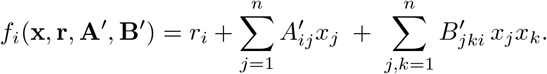

The gradual boundary consists of those parameters **r, A**^*′*^, **B**^*′*^, such that there exists an **x**^⋆^ with

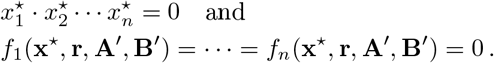

The first condition means that at least one entry of **x**^⋆^ is zero, while the second line implies that **x**^⋆^ is a solution of (3). The Tarski-Seidenberg theorem [54, Theorem 4.17] implies that the gradual boundary is a semialgebraic set (a set defined by polynomial equations and inequalities). While computing all equations and inequalities of a semi-algebraic set is a challenging problem, computing equations is easier, using elimination theory [55, Chapter 3]. The polynomials above generate a polynomial ideal [55]

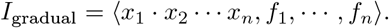

Eliminating **x** from the ideal *I*_gradual_ and substituting back 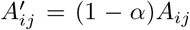 and 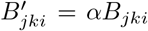 we obtain an ideal *J*_gradual_ generated by polynomials in **r**, *α*, **A, B**, that define the Zariski closure of the gradual boundary (i.e., the smallest algebraic variety – a set defined as the vanishing locus of finitely many polynomials – containing the gradual boundary). We find that *J*_gradual_ is generated by a single polynomial Δ_gradual_ (see SI Sec. 3).

Similarly, for the abrupt boundary we can eliminate **x** from the ideal

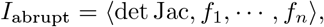

where Jac denotes the Jacobian matrix of the polynomials in (3). Eventually, we obtain the ideal *J*_abrupt_ generated by a single polynomial Δ_abrupt_ in **r**, *α*, **A, B**, that define the Zariski closure of the abrupt boundary (see SI Sec. 3). This polynomial has 17,586 terms, underscoring the complexity of elimination. Recently, methods from numerical algebraic geometry have emerged to tackle elimination ideals [56] which are infeasible to compute using exact computer algebra.

The code to compute Δ_gradual_ and Δ_abrupt_ in `Macaulay2` [57] is in SI Sec. 3.

### Global stability rules out fold bifurcations

A fold bifurcation requires two equilibria to collide and annihilate; if the coexistence equilibrium **x**^⋆^ is globally asymptotically stable on the positive orthant 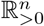, it is the only interior equilibrium and such collisions are precluded. We derive sufficient conditions for global stability using Goh’s Lyapunov function [37, 38, 58]

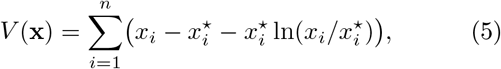

which is non-negative on 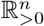 and vanishes only at **x**^⋆^. Using 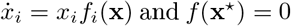and *f* (**x**^⋆^) = 0, together with the path integral *f* (**x**)−*f* (**x**^⋆^) =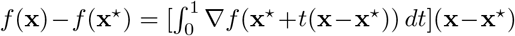, its derivative along trajectories takes the quadratic form

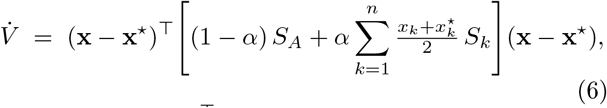

where *S*_*A*_ = (*A*+*A*^T^)/2 and (*S*_*k*_)_*ij*_ = (*B*_*ijk*_ +*B*_*jik*_)/2 (assuming *B*_*ijk*_ = *B*_*ikj*_). Because each 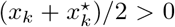 on 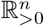, the bracketed matrix is a conic combination of *S*_*A*_ and the slice matrices 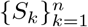. A sufficient condition for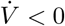 everywhere, and hence for global stability, is therefore that *S*_*A*_ ≺ 0 and *S*_*k*_ ≺ 0 for every *k* = 1, …, *n* - the slice-wise negative-definiteness condition. An equivalent condition arises in the theory of nonlinear complementarity problems via strict monotonicity of *F* = −*f* [59].

The slice-wise condition is strictly stronger than 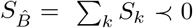, which is enforced by our sampling protocol. The gap between the slice-wise and aggregate conditions delineates the regime in which fold bifurcations can arise (see SI Sec. 5).

## Acknowledgments

P.L.A. acknowledges helpful discussions with E. Alastrue de Asenjo, K. Lee, S. Pawar, S. Kuhen, S. Sheik, M. Tikhonov and T. Wootton.

## Funding

P.B. was supported by DFG, German Research Foundation – Projektnummer 445466444

## Author contributions

P.L.A. designed the study and performed the research, with advice from Z.M. and M.P.; P.L.A. wrote the manuscript, with advice from Z.M. and M.P.; P.L.A., P.B., and A.L. developed the code; P.L.A, Z.M, P.B. and S.A. performed the derivations. All authors contributed to the mathematical derivations, the implementation of the code, edited the manuscript, and discussed and interpreted the results.

## Competing interests

The authors declare no competing interests.

## Notes

### Competing Interest Statement

The authors have declared no competing interest.

