## supplementary information for "Tipping points are typical in ecosystems with higher-order interactions"

#### 1 Model

We consider generalized Lotka–Volterra (GLV) dynamics with higher-order interactions (HOIs):

$$\dot{x}_i = x_i \left[ r_i + (1-\alpha)\sum_{j=1}^{n} A_{ij}\, x_j + \alpha \sum_{j,k=1}^{n} B_{ijk}\, x_j x_k \right], \quad i = 1, \dots, n, \tag{1}$$

where $x_i$ is the abundance of species $i$, $r_i$ its intrinsic growth rate, $A_{ij}$ the pairwise interaction coefficient, and $B_{ijk}$ the three-way higher-order interaction coefficient. The parameter $\alpha \in [0,1]$ controls the relative weight of higher-order versus pairwise terms, recovering the classical GLV model when $\alpha = 0$.

We decompose the growth-rate vector as

$$\mathbf{r} = \mathbf{r}^0 + \delta\,\mathbf{u}, \tag{2}$$

where $\mathbf{r}^0$ is a baseline growth-rate vector that supports a feasible, locally stable coexistence equilibrium, $\mathbf{u}$ is a unit vector specifying the direction of environmental change, and $\delta \geq 0$ its magnitude. By varying $\mathbf{r}$ along fixed directions $\mathbf{u}$, we probe the robustness of coexistence to environmental forcing.

#### 1.1 Feasibility and equilibrium tracking

A coexistence equilibrium $\mathbf{x}^\star \in \mathbb{R}^n_{>0}$ satisfies, for each species,

$$r_i + (1-\alpha)\sum_j A_{ij}\,x_j^\star + \alpha\sum_{j,k} B_{ijk}\,x_j^\star x_k^\star = 0. \tag{3}$$

For $\alpha > 0$ this is a system of $n$ coupled quadratic equations in $\mathbf{x}^\star$, and the inverse problem—determining the set of growth-rate vectors $\mathbf{r}$ for which a positive solution exists—is not analytically tractable in general. We therefore assume a known feasible equilibrium at the baseline $\mathbf{r}^0$ and track the solution branch numerically as $\delta$ varies.

Concretely, when we perturb the growth rate as $\mathbf{r}^0 + \delta\,\mathbf{u}$ (see Eq. (2) in the main text), Eq. (3) becomes

$$H_i(\mathbf{x},\delta) \;=\; (r_i + \delta\,u_i) + (1-\alpha)\sum_j A_{ij}\,x_j + \alpha\sum_{j,k} B_{ijk}\,x_j x_k \;=\; 0, \tag{4}$$

with $H = (H_1, \ldots, H_n)$. Wherever the Jacobian $\frac{\mathrm{d}H}{\mathrm{d}\mathbf{x}}$ is invertible, the implicit function theorem guarantees that $\mathbf{x}^\star(\delta)$ is locally a differentiable function of $\delta$. Implicit differentiation yields the predictor–corrector ODE

$$\frac{\mathrm{d}H}{\mathrm{d}\mathbf{x}}\,\dot{\mathbf{x}} \;+\; \frac{\mathrm{d}H}{\mathrm{d}\delta} \;=\; 0. \tag{5}$$

Together with the initial condition $\mathbf{x}^\star = \mathbf{1}$ at $\delta = 0$, this defines an initial-value problem that we solve using the polynomial homotopy continuation software `HomotopyContinuation.jl` [1, 2].

#### 1.2 Local stability

The local stability of a coexistence equilibrium $\mathbf{x}^\star$ is determined by the Jacobian

$$J_{ij}(\mathbf{x}^\star) = x_i^\star\left[(1-\alpha)\,A_{ij} + \alpha\sum_{k=1}^{n}\big(B_{ijk} + B_{ikj}\big)\,x_k^\star\right] + \underbrace{\delta_{ij}\,f_i(\mathbf{x}^\star)}_{=0}, \tag{6}$$

where $f_i(\mathbf{x}^\star)$ is the per-capita growth rate of species $i$ (the bracketed term in Eq. 1), which vanishes at equilibrium, and $\delta_{ij}$ is the Kronecker delta. The equilibrium is locally stable when all eigenvalues of $J$ have negative real part. Crucially, $J$ depends on $\mathbf{x}^\star$, which itself depends on $\mathbf{r}$ through Eq. (3), establishing the chain of dependence $J\big(\mathbf{x}^\star(\mathbf{r})\big)$. Changes in environmental conditions therefore alter stability, not only through changes in abundances themselves, but also through the state-dependent higher-order terms in the Jacobian.

Equation (6) shows that the community Jacobian factors as

$$J = D(\mathbf{x}^\star)\, J_F, \tag{7}$$

where $D(\mathbf{x}^\star) = \mathrm{diag}(x_1^\star, \ldots, x_n^\star)$ and

$$(J_F)_{ij} \;=\; (1-\alpha)\, A_{ij} \;+\; \alpha \sum_{k=1}^{n} \bigl(B_{ijk} + B_{ikj}\bigr)\, x_k^\star \tag{8}$$

is the Jacobian of the per-capita growth rates. Because the determinant of a product is the product of the determinants,

$$\det J \;=\; \det D(\mathbf{x}^\star)\;\det J_F \;=\; \left(\prod_{i=1}^{n} x_i^\star\right)\,\det J_F. \tag{9}$$

This factorization reveals that $\det J$ can vanish—and hence the equilibrium can undergo a bifurcation—via two distinct mechanisms:

1. **Transcritical bifurcation.** One (or more) equilibrium abundances $x_i^\star \to 0$, so $\prod_i x_i^\star = 0$. A species is smoothly driven to extinction.
2. **Fold (saddle–node) bifurcation.** All abundances remain positive, but $\det J_F = 0$. The per-capita Jacobian $J_F$ becomes singular while every species is still present.

Note that $J_F$ coincides with the Jacobian $\frac{\mathrm{d}H}{\mathrm{d}\mathbf{x}}$ of the homotopy system (4). At a fold, therefore, the implicit function theorem fails: $\frac{\mathrm{d}H}{\mathrm{d}\mathbf{x}}$ is no longer invertible, so the equilibrium branch $\mathbf{x}^\star(\delta)$ ceases to be a smooth, real-valued function of $\delta$. The continuation path cannot be extended further as a real solution, and the equilibrium disappears abruptly. The next section describes how we detect and distinguish these two boundary types in practice.

#### 2 Boundary detection with homotopy continuation

For each perturbation direction $\mathbf{u}$, we track the coexistence equilibrium from $\delta = 0$ to a pre-set upper limit $\delta_{\max}$ using coefficient-parameter homotopy continuation (see Algorithm 1). The tracker advances a homotopy parameter $t \in [1, 0]$, where $t = 1$ corresponds to $\delta = 0$ and $t = 0$ to $\delta = \delta_{\max}$, smoothly deforming the model parameters along the ray $\mathbf{r}_0 + \delta\,\mathbf{u}$ and carrying the equilibrium branch $\mathbf{x}(t)$ along with it. At each continuation step we monitor the tracked solution for three types of boundary events: *gradual boundary*, where a species abundance $x_i$ crosses zero; *stability loss*, where the real part of the leading eigenvalue $\lambda_{\max}$ of the Jacobian crosses zero; and *abrupt boundaries*, detected when the tracker itself fails to converge, indicating that the equilibrium branch has ceased to exist via a saddle-node bifurcation (see Fig. S1). When a feasibility or stability event is detected between two consecutive continuation steps, a bisection procedure refines the critical parameter value to within tolerance (red band in Fig. S1). The three boundary types and the no-boundary case are defined precisely as follows.

1. **Gradual boundary** (`transcritical`): The minimum component of the real part of the tracked equilibrium drops below $x_{tol} = 10^{-12}$. This signals that at least one species has been driven to zero abundance, corresponding to a transcritical bifurcation.

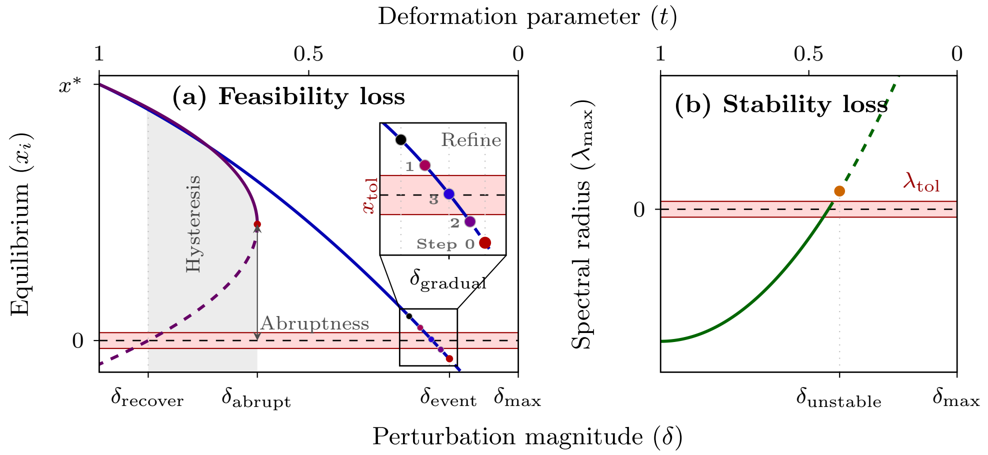


**Fig. S1** Schematic of the three boundary types detected by the homotopy continuation algorithm as a function of perturbation magnitude $\delta$ (equivalently, deformation parameter $t = 1 - \delta/\delta_{max}$). (a) Feasibility loss. Two mechanisms are shown: a gradual boundary (blue), where a species equilibrium $x_i$ smoothly declines and crosses the feasibility threshold $x_{tol}$ at $\delta_{gradual}$; and an abrupt boundary (violet), where a saddle-node bifurcation annihilates the stable equilibrium at $\delta_{abrupt}$, causing a discontinuous drop in abundance. The vertical distance from the fold point to zero quantifies the abruptness of the transition. The shaded region between $\delta_{recover}$ and $\delta_{abrupt}$ marks the hysteresis zone, where the system cannot recover by simply reversing the perturbation. (b) Stability loss. The leading eigenvalue $\lambda_{max}$ increases with $\delta$ and crosses the stability threshold $-\lambda_{tol}$ at $\delta_{unstable}$, signalling loss of dynamical stability before any species becomes infeasible.

2. **Unstable boundary** (`hopf`): The equilibrium remains feasible (all components real and positive) but loses local stability. Stability is monitored by computing the maximum real part $\lambda_{max}$ of the eigenvalues of the community Jacobian $J = D(\mathbf{x}^\star)J_F$, where $D(\mathbf{x})$ is a diagonal matrix with the vector $\mathbf{x}$ in the diagonal, and $J_F$ is the Jacobian of the per-capita growth rates (Eq. 10). The equilibrium is declared unstable when $\lambda_{max} > 10^{-9}$.
3. **Abrupt boundary** (`fold`): The tracker fails. This signals a fold (saddle–node) bifurcation where the real equilibrium branch ceases to exist. The termination occurs because the Jacobian $J_F$ of the per-capita growth rates (Eq. 10) becomes singular as the fold is approached: at a fold bifurcation, two real solution branches (one stable, one unstable) collide and annihilate, so the matrix $\frac{\mathrm{d}H}{\mathrm{d}\mathbf{x}}$ in Eq. 5 ceases to be invertible. Since the implicit function theorem requires invertibility of this Jacobian to guarantee that $\mathbf{x}(\delta)$ remains a smooth, real-valued function of $\delta$, the theorem's hypotheses fail at the fold. Consequently, the continuation path cannot be extended further as a real solution branch: the equilibrium leaves the real line.
4. **No boundary** (`success`): The equilibrium remains feasible and stable up to $\delta_{max}$.

When a gradual or unstable boundary event is detected between consecutive homotopy-continuation steps at path parameters $t_{prev}$ (last safe state, black marker in Fig. S1(a), inset) and $t_{cur}$ (first event state, red, Step 0, marker in the inset), the event location is refined by re-tracking on progressively smaller $t$-intervals; a bisection algorithm that finds solution via homotopy continuation rather than via Newton method.

Concretely, the tracker is re-initialised at the state $(\mathbf{x}_{cur}, t_{cur})$ with the goal of stepping back toward $t_{prev}$. The adaptive step-size control is tightened: the maximum

**Algorithm 1** Boundary detection via homotopy continuation

**Require:** Equilibrium $\mathbf{x}^*$, baseline parameters $\mathbf{r}_0$, direction $\mathbf{u}$ ($\|\mathbf{u}\| = 1$), max perturbation $\delta_{\max}$, extinction threshold $x_{tol}$, stability tolerance $\lambda_{\mathrm{tol}}$, parameter tolerance $\tau$
**Ensure:** Boundary type, critical perturbation $\Delta\mathbf{r}_c$, critical equilibrium $\mathbf{x}_c$

Initialize homotopy from $\mathbf{p}_0 = \mathbf{0}$ to $\mathbf{p}_1 = \delta_{\max}\,\mathbf{u}$, starting at $\mathbf{x}^*$
**while** continuation path is active **do**
  Advance tracker one step $\rightarrow$ update $\mathbf{x}_{\mathrm{cur}}$, $t$
  **if** any $x_{\mathrm{cur},i} < -x_{tol}$ **then** ▷ Transcritical
    Refine $t$ by bisection until $|x_i| \leq x_{tol}$ or $\Delta t < \tau$
    **return** NEGATIVE, $(1-t)\,\mathbf{p}_1$, $\mathbf{x}_{\mathrm{cur}}$
  **end if**
  **if** $\lambda_{\max}$ of community matrix crossed $\lambda_{\mathrm{tol}}$ **then** ▷ Hopf
    Refine $t$ by bisection until $|\lambda_{\max} - \lambda_{\mathrm{tol}}| \leq \lambda_{\mathrm{tol}}$ or $\Delta t < \tau$
    **return** UNSTABLE, $(1-t)\,\mathbf{p}_1$, $\mathbf{x}_{\mathrm{cur}}$
  **end if**
  **if** tracker diverged **then return** FOLD, $(1-t)\,\mathbf{p}_1$, $\mathbf{x}_{\mathrm{cur}}$ ▷ Fold
  **end if**
**end while**
**return** SUCCESS, $\mathbf{p}_1$, $\mathbf{x}_{\mathrm{cur}}$ ▷ No boundary found

step size is set to $\Delta t/2$ where $\Delta t = |t_{\mathrm{prev}} - t_{\mathrm{cur}}|$, and the maximum number of steps is set to $\lceil 1/\Delta t \rceil$.

At each refinement step, the tracker advances and the relevant event indicator is re-evaluated:

- **Negative event** (`find_zero`): the indicator is the abundance $x_i$ of the species that crossed zero. If $|x_i| \leq x_{tol}$, the current $t$ is accepted. If $x_i$ overshoots (changes sign relative to the initial direction), the function *recurses* on the sub-interval $[t'_{\mathrm{prev}}, t'_{\mathrm{cur}}]$ bracketed by the last two tracker states.
- **Unstable event** (`find_stability`): the indicator is $\lambda_{\max} - \lambda_{\mathrm{tol}}$. If $|\lambda_{\max} - \lambda_{\mathrm{tol}}| \leq \lambda_{\mathrm{tol}}$, $t$ is accepted. If $\lambda_{\max}$ crosses back below the threshold, the function recurses on the new sub-interval.

Because a transcritical bifurcation simultaneously drives $x_i \rightarrow 0$ and generically pushes $\lambda_{\max}$ through zero (the zero-abundance eigenvector enters the community matrix), the two events can coincide along a perturbation ray. Concretely, if at homotopy parameter $t^*$ both $x_i(t^*) \leq x_{tol}$ and $\lambda_{\max}(t^*) \geq \lambda_{\mathrm{tol}}$, the event is classified as `negative` (feasibility loss via transcritical bifurcation), not `unstable`.

Refinement terminates when either (i) the indicator is within tolerance, (ii) the tracker step size $\Delta t$ collapses to zero, or (iii) the tracker can no longer reduce the interval (recursive base case). The final homotopy parameter $t_{\mathrm{end}}$ determines the critical perturbation via $\Delta\mathbf{r}_c = (1-, t_{\mathrm{end}})\,\mathbf{p}_{\mathrm{target}}$ and $\delta_c = \|\Delta\mathbf{r}_c\|$.

### 3 Computing the feasibility boundaries analytically for $n = 2$

We discuss how to compute the zero- and fold/complex-boundary analytically. For this we substitute in (3) $A'_{ij} = (1-\alpha)A_{ij}$ and $B'_{jki} = \alpha B_{jki}$, and obtain polynomials

$$f_i(\mathbf{x}, \mathbf{r}, \mathbf{A}', \mathbf{B}') = r_i + \sum_{j=1}^{n} A'_{ij} x_j \; + \sum_{j,k=1}^{n} B'_{jki}\, x_j x_k.$$

The zero-boundary consist of those parameters $\mathbf{r}, \mathbf{A}', \mathbf{B}'$, such that there exists an $\mathbf{x}^\star$ with

$$\begin{aligned} &x_1^\star \cdot x_2^\star \cdots x_n^\star = 0 \quad \text{and} \\ &f_1(\mathbf{x}^\star, \mathbf{r}, \mathbf{A}', \mathbf{B}') = \cdots = f_n(\mathbf{x}^\star, \mathbf{r}, \mathbf{A}', \mathbf{B}') = 0 \end{aligned}$$

The first condition means that at least one entry of $\mathbf{x}^\star$ is zero, while the second line implies that $\mathbf{x}^\star$ is a solution of (3). These polynomials generate a *polynomial ideal* [3]

$$I_{\text{zero}} = \langle x_1 \cdot x_2 \cdots x_n, f_1, \cdots, f_n \rangle.$$

Using the theory of elimination (see [3, Chapter 3]), we can eliminate $\mathbf{x}$ from this ideal. Then, we substitute back $A'_{ij} = (1-\alpha)A_{ij}$ and $B'_{jki} = \alpha B_{jki}$, and obtain an ideal $J_{\text{zero}}$ generated by polynomials in $\mathbf{r}, \alpha, \mathbf{A}, \mathbf{B}$, that define the Zariski-closure of the gradual boundary. Elimination can be done on a computer, for instance, using the computer algebra system `Macaulay2` We find that the zero-boundary is the zero set of a single polynomial; i.e., $J_{\text{zero}}$ is a principal ideal generated by

$$\Delta_{\text{zero}} = \alpha^2 \cdot P \cdot Q,$$

where

$$\begin{aligned} Q = {} & r_1^2 \alpha B_{222}^2 - 2 r_1 r_2 \alpha B_{122} B_{222} - r_1 \alpha^2 A_{12} A_{22} B_{222} \\ & + r_1 \alpha^2 A_{22}^2 B_{122} + 2 r_1 \alpha A_{12} A_{22} B_{222} - 2 r_1 \alpha A_{22}^2 B_{122} \\ & - r_1 A_{12} A_{22} B_{222} + r_1 A_{22}^2 B_{122} + r_2^2 \alpha B_{122}^2 + r_2 \alpha^2 A_{12}^2 B_{222} \\ & - r_2 \alpha^2 A_{12} A_{22} B_{122} - 2 r_2 \alpha A_{12}^2 B_{222} + 2 r_2 \alpha A_{12} A_{22} B_{122} \\ & + r_2 A_{12}^2 B_{222} - r_2 A_{12} A_{22} B_{122}; \\ P = {} & r_1^2 \alpha B_{211}^2 - 2 r_1 r_2 \alpha B_{111} B_{211} - r_1 \alpha^2 A_{11} A_{21} B_{211} \\ & + r_1 \alpha^2 A_{21}^2 B_{111} + 2 r_1 \alpha A_{11} A_{21} B_{211} - 2 r_1 \alpha A_{21}^2 B_{111} \\ & - r_1 A_{11} A_{21} B_{211} + r_1 A_{21}^2 B_{111} + r_2^2 \alpha B_{111}^2 + r_2 \alpha^2 A_{11}^2 B_{211} \\ & - r_2 \alpha^2 A_{11} A_{21} B_{111} - 2 r_2 \alpha A_{11}^2 B_{211} + 2 r_2 \alpha A_{11} A_{21} B_{111} \\ & + r_2 A_{11}^2 B_{211} - r_2 A_{11} A_{21} B_{111}. \end{aligned}$$

Similarly, for the complex boundary we can eliminate $\mathbf{x}$ from the ideal

$$I_{\text{complex}} = \langle \det J_F, f_1, \cdots, f_n \rangle,$$

where $J_F$ denotes the Jacobian matrix of the per-capita growth rates, i.e.,

$$(J_F)_{ij} = \frac{\partial f_i}{\partial x_j} = (1-\alpha)A_{ij} + \alpha \sum_k \big(B_{ijk} + B_{ikj}\big)x_k. \tag{10}$$

The eliminant is called a *tact invariant* in algebraic geometry. It defines the locus of pairs of quadrics in the plane that meet tangentially [4]. Eventually, we obtain the ideal $I_{\text{complex}}$ generated by polynomials in $\mathbf{r}, \alpha, \mathbf{A}, \mathbf{B}$, that define the Zariski-closure of the complex-boundary. Again, $J_{\text{complex}}$ is generated by the polynomial

$$\begin{aligned}
\Delta_{\text{complex}} = \alpha^4 \cdot \big(&256 r_1^4 \alpha^4 B_{111}^2 B_{211}^2 B_{222}^4 \\
&- 128 r_1^4 \alpha^4 B_{111}^2 B_{211} B_{212}^2 B_{222}^3 \\
&+ 16 r_1^4 \alpha^4 B_{111}^2 B_{212}^4 B_{222}^2 \\
&- 256 r_1^4 \alpha^4 B_{111} B_{112} B_{211}^2 B_{212} B_{222}^3 \\
&+ 128 r_1^4 \alpha^4 B_{111} B_{112} B_{211} B_{212}^3 B_{222}^2 \\
&- 16 r_1^4 \alpha^4 B_{111} B_{112} B_{212}^5 B_{222} - \cdots \\
&\quad \cdots \\
&- 8 A_{12}^2 A_{21}^4 A_{22}^2 B_{111}^2 B_{122}^2 - 2 A_{12}^2 A_{21}^4 A_{22}^2 B_{111} B_{112}^2 B_{122} \\
&+ A_{12}^2 A_{21}^4 A_{22}^2 B_{112}^4 + 8 A_{12}^2 A_{21}^3 A_{22}^3 B_{111}^2 B_{112} B_{122} \\
&- 2 A_{12}^2 A_{21}^3 A_{22}^3 B_{111} B_{112}^3 - 4 A_{12}^2 A_{21}^2 A_{22}^4 B_{111}^3 B_{122} \\
&+ A_{12}^2 A_{21}^2 A_{22}^4 B_{111}^2 B_{112}^2\big).
\end{aligned}$$

This polynomial has 17.586 terms, which underscores the complexity of elimination. For bigger $n$ computing elimination ideals quickly becomes infeasible. Recently, methods from numerical algebraic geometry have emerged to tackle elimination ideals [5] which are infeasible to compute using exact computer algebra.

### 4 Parameter sampling protocol: construction, scaling, and constraints

Our simulations pursue two goals that pull in opposite directions. On one hand, we want communities drawn from a broad random ensemble so that results reflect the generic, typical behavior of ecosystems with higher-order interactions rather than the idiosyncrasies of any particular parameter choice. On the other hand, we want a controlled experimental design in which the quantities of interest—HOI strength $\alpha$—is the sole unconfounded variable, so that differences in boundary composition can be attributed causally to the nonlinearity or sign bias of interactions rather than to a shifting reference state, a changing interaction magnitude, or an unstable baseline. We reconcile these goals by imposing structure only where causal attribution demands it, and leaving everything else to random variation.

Concretely, we draw interaction coefficients independently from broad distributions, ensuring ensemble diversity, but we then apply three targeted constraints: (i) a planted equilibrium, so that all communities share the same baseline coexistence state $\mathbf{x}^\star = \mathbf{1}$ at $\mathbf{r}^0 = \mathbf{1}$ regardless of $\alpha$, $\mu_A$, and $\mu_B$; and (ii) a spectral normalisation that guarantees local stability at the baseline for all $\alpha$. We describe these steps in turn.

Off-diagonal entries are drawn independently:

$$\tilde{A}_{ij} \sim \mathcal{N}(\mu_A,\ \tfrac{1}{n}), \qquad \tilde{B}_{ijk} \sim \mathcal{N}(\mu_B,\ \tfrac{1}{n^2}), \tag{11}$$

where $\mu_A$ and $\mu_B$ control the mean sign of pairwise and higher-order interactions, respectively. In our sign convention, $\mu_A > 0$ and $\mu_B > 0$ correspond to net facilitation, while $\mu_A < 0$ and $\mu_B < 0$ correspond to net competition. The variance scalings ensure that both the pairwise and aggregated HOI contributions to the per-capita growth rate remain $O(1)$ as community size $n$ grows: the row sum $\sum_j A_{ij}x_j$ involves $n$ terms each of variance $1/n$, and the slice sum $\sum_{j,k} B_{ijk}x_jx_k$ involves $n^2$ terms each of variance $1/n^2$. The construction below applies uniformly for any choice of $(\mu_A, \mu_B)$; the main text studies three regimes varying $\mu_B$ at $\mu_A = 0$, and Appendix B extends this to variable $\mu_A$.

Next, we aim to construct $A$ and $B$ from the random draws $\tilde{A}$, $\tilde{B}$ so that $\mathbf{x}^\star = \mathbf{1}$ is a locally stable equilibrium for all $\alpha$. The Jacobian (6) evaluated at $\mathbf{x}^\star = \mathbf{1}$ takes the form

$$J(\alpha) = (1-\alpha)\,A + \alpha\,\hat{B}, \tag{12}$$

where $\hat{B}_{ij} = \sum_k (B_{ijk} + B_{ikj})$. Define the symmetric parts of the pairwise and aggregated HOI matrices as

$$S_A = \tfrac{1}{2}(A + A^\top), \qquad (S_{\hat{B}})_{ij} = \tfrac{1}{4}\sum_k \big(B_{ijk} + B_{ikj} + B_{jik} + B_{jki}\big). \tag{13}$$

A sufficient condition for Hurwitz stability is that the symmetrised combination is negative definite: $(1-\alpha)\,S_A + \alpha\,S_{\hat{B}} \prec 0$. Since this is a convex combination with non-negative weights, it suffices that both sectors be separately negative definite:

$$S_A \prec 0 \qquad \text{and} \qquad S_{\hat{B}} \prec 0. \tag{14}$$

***Step 1: zero-sum diagonals.***

We set the diagonal entries of $\tilde{A}$ and $\tilde{B}$ to enforce zero row and slice sums:

$$\tilde{A}_{ii} = -\sum_{j \neq i} \tilde{A}_{ij}, \qquad \tilde{B}_{iii} = -\sum_{(j,k)\neq(i,i)} \tilde{B}_{ijk}, \tag{15}$$

so that $\sum_j \tilde{A}_{ij} = 0$ and $\sum_{j,k} \tilde{B}_{ijk} = 0$ for every species $i$. Off-diagonal entries are unchanged.

***Step 2: rescaling and self-regulation.***

The raw matrices $\tilde{A}$ and $\tilde{B}$ will not generally satisfy (14). A natural approach is to subtract a diagonal shift, giving interaction matrices $\tilde{A}_{ij} - \delta_A\,\delta_{ij}$ and $\tilde{B}_{ijk} - \delta_B\,\delta_{ijk}$ for sufficiently large $\delta_A, \delta_B > 0$. However, this changes the row and slice sums from zero to $-\delta_A$ and $-\delta_B$ respectively, so the equilibrium condition at $\mathbf{x} = \mathbf{1}$ would break. We resolve this by observing that, for any $\delta_A > 0$,

$$\tilde{A}_{ij} - \delta_A\,\delta_{ij} \;=\; \delta_A \left( \frac{\tilde{A}_{ij}}{\delta_A} - \delta_{ij} \right). \tag{16}$$

Since multiplying a matrix by a positive scalar preserves the sign of every eigenvalue, the symmetric part of the matrix $\tilde{A} - \delta_A\,I$ is negative definite if and only if that of $\tilde{A}/\delta_A - I$ is. The same factorisation applies slice-wise to the HOI tensor. The final interaction matrices are therefore

$$A_{ij} = \frac{\tilde{A}_{ij}}{\delta_A} - \delta_{ij}, \qquad B_{ijk} = \frac{\tilde{B}_{ijk}}{\delta_B} - \delta_{ijk}, \tag{17}$$

where $\delta_{ij}$ is the Kronecker delta, $\delta_{ijk} = 1$ iff $i = j = k$, and the spectral thresholds $\delta_A, \delta_B > 0$ are determined below.

This construction simultaneously achieves both goals. The zero-sum properties are preserved under uniform rescaling ($\sum_j A_{ij} = 0/\delta_A - 1 = -1$ and $\sum_{j,k} B_{ijk} = 0/\delta_B - 1 = -1$), so the equilibrium condition at $\mathbf{x}^\star = \mathbf{1}$ with $r_i = 1$ reads

$$1 + (1-\alpha)(-1) + \alpha(-1) = 0,$$

which holds identically for all $\alpha \in [0, 1]$, independent of $\delta_A$, $\delta_B$, $\mu_A$, and $\mu_B$. At the same time, the stability conditions (14) reduce to constraints on the spectral thresholds, which we now determine.

***Choosing the spectral thresholds.***

The spectral thresholds $\delta_A$ and $\delta_B$ must each absorb two contributions: the bulk fluctuations of the random matrix and any mean-field spike induced by a nonzero interaction mean. We handle both sectors in parallel, decomposing each symmetric matrix into its deterministic mean part and a centred (zero-mean) fluctuation:

$$S_A = \mathbb{E}[S_A] + \tilde{S}_A, \qquad S_{\hat{B}} = \mathbb{E}[S_{\hat{B}}] + \tilde{S}_{\hat{B}}, \tag{18}$$

where $\tilde{S}_A$ and $\tilde{S}_{\hat{B}}$ denote the centred parts. In Appendix A we show that both mean matrices share the same centring-matrix structure:

$$\mathbb{E}[S_A] = -n\mu_A\left(I - \frac{\mathbf{1}\mathbf{1}^\top}{n}\right), \qquad \mathbb{E}[S_{\hat{B}}] = -n^2\mu_B\left(I - \frac{\mathbf{1}\mathbf{1}^\top}{n}\right), \tag{19}$$

with spectra $n\mu_A$ (mult. $n-1$), 0 (eigenvector $\mathbf{1}$) and $n^2\mu_B$ (mult. $n-1$), 0 (eigenvector $\mathbf{1}$) respectively. The $O(n)$ versus $O(n^2)$ prefactors reflect the number of terms aggregated in each sector: $n$ pairwise terms per row versus $n^2$ HOI terms per slice. In both cases, the sign of the mean determines whether the deterministic spike stabilises or destabilises the symmetric interaction matrix, and the structure is identical across sectors: facilitative means ($\mu_A, \mu_B > 0$) produce a negative spike and are stabilising, while competitive means ($\mu_A, \mu_B < 0$) produce a positive spike and are destabilising. Figure S2 illustrates this for the HOI sector; the pairwise sector is qualitatively identical with $O(n)$ rather than $O(n^2)$ spike magnitude.

Because the three regimes are structurally identical across sectors, we describe them once and apply the prescription to both $\delta_A$ and $\delta_B$. Let $S$ denote either $S_A$ or $S_{\hat{B}}$, let $\tilde{S}$ denote its centred part, and let $\mu \in \{\mu_A, \mu_B\}$ denote the corresponding interaction mean. From the factorisation (16), stability requires $\delta > \lambda_{\max}(S)$; the regime-dependent prescription is as follows.

*Facilitative regime ($\mu > 0$; stabilising spike).* The $(n-1)$-fold eigenvalue of $\mathbb{E}[S]$ is negative, pushing the bulk of the spectrum of $S$ deep into the negative half-line, while $\lambda_{\max}(\mathbb{E}[S]) = 0$ along the $\mathbf{1}$ direction. Any threshold $\delta > \lambda_{\max}(\tilde{S})$ suffices for negative definiteness. To maintain a uniform construction across regimes, we set the threshold

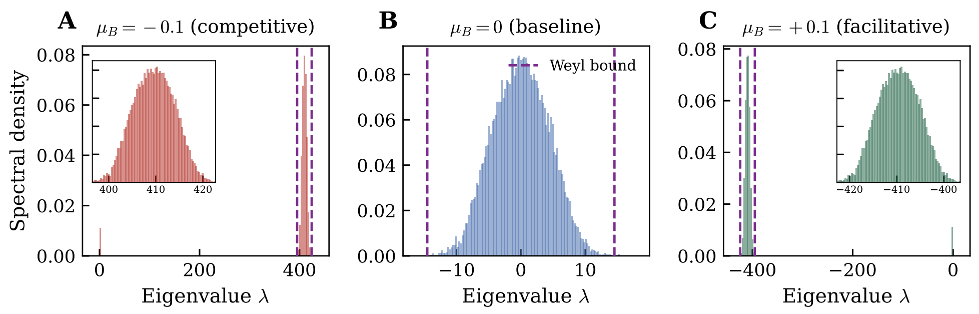


**Fig. S2** Spectral density of the symmetric HOI aggregation matrix $S_{\hat{B}}$ for $n = 32$ across the three $\mu_B$ regimes. In the baseline ($\mu_B = 0$), the spectrum is centred at zero. In the facilitative ($\mu_B > 0$) and competitive ($\mu_B < 0$) regimes, the mean-field eigenvalues of $\mathbb{E}[S_{\hat{B}}]$ shift the $(n-1)$-fold bulk away from the origin (to the left under facilitation, to the right under competition), while the top eigenvalue (near the **1** direction) remains near the fluctuation edge. The pairwise matrix $S_A$ exhibits analogous behaviour across $\mu_A$ regimes, with the spike scaled by $n\mu_A$ rather than $n^2\mu_B$.

using the fluctuation matrix:

$$\delta_A = \lambda_{\max}(\tilde{S}_A), \qquad \delta_B = \lambda_{\max}(\tilde{S}_{\hat{B}}),$$

both of which inherit $O(\sqrt{\log n})$ scaling (Appendix B).

*Baseline ($\mu = 0$).* The mean matrix vanishes: $\mathbb{E}[S] = 0$, so $\tilde{S} = S$ and the prescription above reduces to $\delta_A = \lambda_{\max}(S_A)$ and $\delta_B = \lambda_{\max}(S_{\hat{B}})$, with the same $O(\sqrt{\log n})$ asymptotics.

*Competitive regime ($\mu < 0$; destabilising spike).* The $(n-1)$-fold eigenvalue of $\mathbb{E}[S]$ flips positive, producing a spectral spike that dominates the top eigenvalue of the full matrix: $\lambda_{\max}(S_A) \sim n|\mu_A|$ and $\lambda_{\max}(S_{\hat{B}}) \sim n^2|\mu_B|$. The threshold must absorb the spike directly:

$$\delta_A = \lambda_{\max}(S_A) = O(n), \qquad \delta_B = \lambda_{\max}(S_{\hat{B}}) = O(n^2).$$

Ecologically, this reflects a basic constraint: predominantly competitive communities require commensurately strong self-regulation to prevent species from suppressing one another to extinction [6, 7], with the required self-regulation strength scaling with community size. The asymmetry between the pairwise ($O(n)$) and HOI ($O(n^2)$) scalings reflects the larger combinatorial footprint of three-way interactions.

###### *Results across $(\mu_A, \mu_B)$ combinations.*

In the main text, we presented the HOI mean $\mu_B$ while fixing $\mu_A = 0$. To verify that the resulting fold-prevalence patterns do not depend on the particular choice of $\mu_A$, we repeated the full boundary-detection protocol across a $3 \times 3$ grid of interaction means $(\mu_A, \mu_B) \in \{-0.1, 0, 0.1\}$, keeping all other aspects of the sampling protocol unchanged (Fig. S3). The qualitative signatures identified in the main text survive across all nine combinations: fold boundaries become increasingly prevalent as $\alpha$ grows; facilitative HOIs ($\mu_B > 0$) produce the steepest and most complete transition to the fold-dominated regime; and the effect of community size reverses sign across $\mu_B$ (folds become rarer with $n$ under competitive or neutral HOIs, but more common under facilitative HOIs). Shifting $\mu_A$ modulates the $\alpha$-location at which folds start to dominate but does not alter these qualitative trends, confirming that the main-text

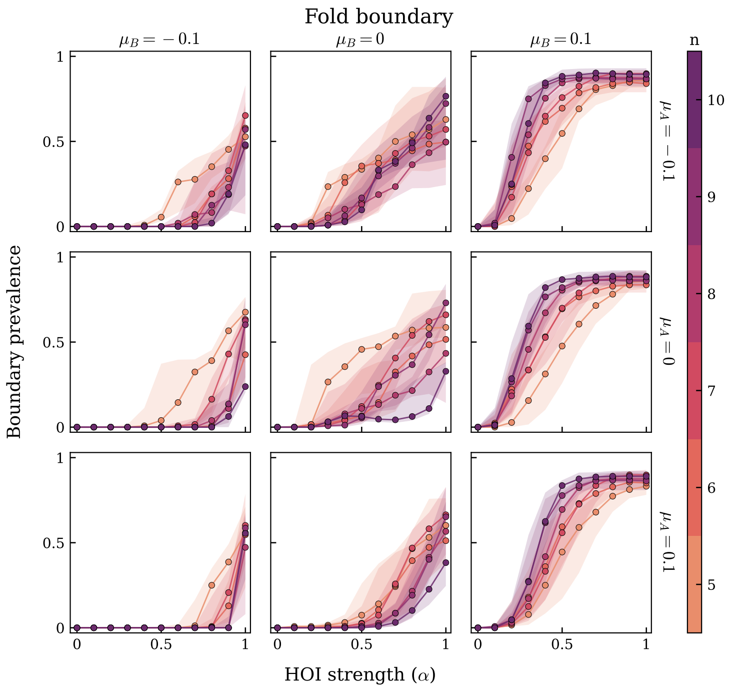


**Fig. S3 Fold-boundary prevalence across the $(\mu_A, \mu_B)$ plane.** Each panel shows the fraction of perturbation directions terminating at a fold (abrupt) boundary as a function of HOI strength $\alpha$, for community sizes $n = 5$–10 (colours). Columns vary the mean higher-order interaction $\mu_B \in \{-0.1, 0, 0.1\}$ (competitive, mixed, and facilitative HOI regimes); rows vary the mean pairwise interaction $\mu_A \in \{-0.1, 0, 0.1\}$ (competitive, mixed, and facilitative pairwise regimes). Points are medians across 50 community replicates at each $(n, \alpha)$ condition; shaded bands span the interquartile range. The central row ($\mu_A = 0$, all three values of $\mu_B$) reproduces Fig. 3 of the main text. The six surrounding panels confirm that the three qualitative patterns established in the main text—(i) fold prevalence rises monotonically with $\alpha$, (ii) facilitative HOIs ($\mu_B > 0$) produce the earliest and most complete transition to the fold-dominated regime, and (iii) the effect of community size on fold prevalence reverses sign across $\mu_B$ (diversity suppresses folds when $\mu_B \leq 0$ but amplifies them when $\mu_B > 0$)—are robust to the mean sign of pairwise interactions. Variation in $\mu_A$ modulates the quantitative onset of fold dominance but does not alter these qualitative trends.

results reflect generic features of the HOI sign structure rather than artefacts of the $\mu_A = 0$ baseline.

#### 4.1 Purely competitive systems

Fold bifurcations have traditionally been associated with positive-feedback architectures—mutualistic loops, Allee effects, cooperative nonlinearities—in which a subset of interactions must be net-facilitative for a saddle–node to arise [8]. In our sampling protocol, by contrast, off-diagonal entries of $A$ and $B$ are drawn from symmetric distributions and therefore contain both positive and negative coefficients even in the $\mu_A = \mu_B = 0$ regime. A natural objection is that the folds we observe might simply reflect the presence of this positive minority: remove the positive entries and the folds would disappear. This section tests that hypothesis directly.

We examine a parameterization in which every interaction coefficient is strictly non-positive by construction. Off-diagonal and diagonal entries alike are drawn from a half-normal distribution and negated:

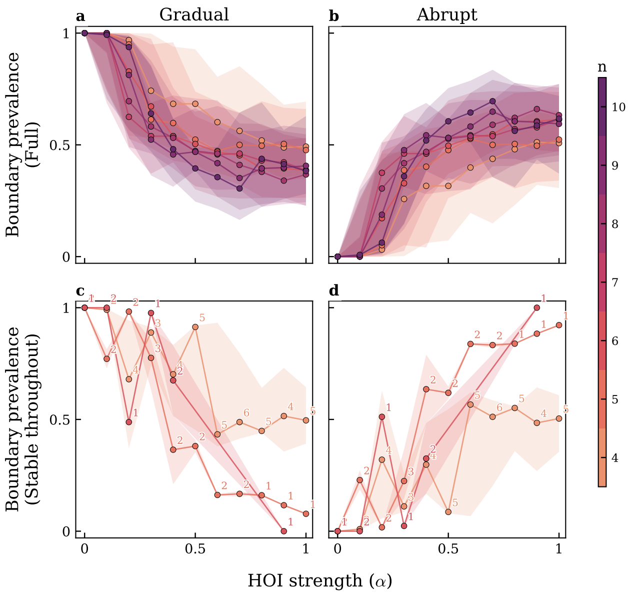


**Fig. S4 Boundary-type prevalence under the purely competitive parameterization.** Each panel shows the fraction of perturbation directions terminating at a given boundary type as a function of HOI strength $\alpha$, for community sizes $n = 4$–$10$ (colours). Columns correspond to abrupt (fold) and gradual (negative-transcritical) boundaries. **Top row:** full bank, containing every replicate with a feasible equilibrium at $\alpha = 0$, including those that later lose stability as $\alpha$ increases. **Bottom row:** stable-throughout subset, restricted to replicates whose coexistence equilibrium $\mathbf{x}^\star = \mathbf{1}$ remains locally stable across the entire $\alpha$ grid, matching the stability condition imposed by construction in the main-text protocol. Points are medians across replicates; bars span the interquartile range. Numerals in the bottom row report the replicate count contributing to each point. In both subsets, fold prevalence rises monotonically with $\alpha$ while gradual-boundary prevalence falls, matching the qualitative pattern of the main text and of Fig. S3. The signature therefore persists in a sampling protocol that removes both the zero-sum-diagonal construction and the spectral rescaling used in the main text, retaining only a planted equilibrium for a purely competitive community.

$$A_{ij} = -\big|\tilde{A}_{ij}\big|, \quad \tilde{A}_{ij} \sim \mathcal{N}\big(0,\, \sigma_A^2/n\big); \qquad B_{ijk} = -\big|\tilde{B}_{ijk}\big|, \quad \tilde{B}_{ijk} \sim \mathcal{N}\big(0,\, \sigma_B^2/n^2\big), \tag{20}$$

with $\sigma_A = \sigma_B = 1$ so that per-capita contributions remain $O(1)$ in $n$. The community is therefore purely competitive: every pairwise and higher-order interaction has its sign fixed a priori, and the resulting $A$ and $B$ bear no spectral-rescaling structure. We instead set the growth rates to the equilibrium condition at $\mathbf{x}^\star = \mathbf{1}$:

$$r_i(\alpha) = -\Big[(1-\alpha)\sum_j A_{ij} + \alpha \sum_{j,k} B_{jki}\Big], \tag{21}$$

Crucially, nothing guarantees that this equilibrium is locally stable: the Jacobian spectrum at $\mathbf{x}^\star$ is determined entirely by the random draw, and stability may fail at some or all values of $\alpha$. This protocol therefore strips away both the zero-sum construction and the $\alpha$-uniform stability guarantee, leaving only the bare planted equilibrium and the raw competitive-sign constraint.

Because stability is no longer enforced by construction, we partition the resulting bank into two subsets (Fig. S4). The *full* bank retains every replicate with a feasible equilibrium at $\alpha = 0$, including those that lose stability as $\alpha$ grows, or that were not stable to begin with. The *stable-throughout* subset retains only those replicates whose coexistence equilibrium is locally stable across the entire $\alpha$ grid, matching the stability condition that the main-text construction imposes by design. Comparing the two subsets isolates the contribution of the $\alpha$-uniform stability guarantee from the other elements of the sampling protocol.

In both subsets, fold-boundary prevalence rises monotonically with $\alpha$ while the prevalence of gradual (negative-transcritical) boundaries declines—the same qualitative signature established in the main text and reinforced across the $(\mu_A, \mu_B)$ grid of Fig. S3. The effect is quantitatively sharper in the stable-throughout subset, but its direction, monotonicity, and dependence on community size are preserved in the unrestricted bank as well. We conclude that the fold-dominance signature is not a consequence of the zero-sum-diagonal construction, of the particular spectral rescaling, or of the flexibility to sample interactions of either sign: it persists in a protocol whose only structural element is a planted equilibrium for a community of purely competitive interactions.

### 5 Global stability rules out fold bifurcations

The results of the previous section show that the statistical properties of the interaction tensors—their means, variances, and the relative weight of higher-order terms—exert a decisive influence on the prevalence of fold bifurcations at feasibility boundaries. In some regimes folds are virtually guaranteed; in others they are rare. This raises a natural question: under what conditions on the interaction tensors $A$ and $B$ can fold bifurcations be ruled out entirely? Here we derive a sufficient condition directly on $A$ and $B$ that guarantees global asymptotic stability of the coexistence equilibrium on the positive orthant, and thereby precludes fold bifurcations at the feasibility boundary.

A fold (saddle–node) bifurcation is, by definition, the collision and annihilation of two equilibrium branches as a parameter is varied [9]. At the fold point, two real equilibria merge; on either side of the fold, the local count of real equilibria differs by two. Consequently, a necessary condition for the existence of a fold bifurcation in parameter space is that the system admit more than one non-negative equilibrium for some values of the parameters.

#### 5.1 Global stability implies uniqueness

A standard condition that implies uniqueness of the coexistence equilibrium is global asymptotic stability on the positive orthant $\mathbb{R}^n_{>0}$. If every trajectory initiated in $\mathbb{R}^n_{>0}$ converges to a single equilibrium $\mathbf{x}^\star$, then $\mathbf{x}^\star$ is necessarily the only equilibrium in $\mathbb{R}^n_{>0}$: any other interior equilibrium would itself be a fixed point of the dynamics and hence could not converge to $\mathbf{x}^\star$, contradicting global convergence. Combined with the observation above, global asymptotic stability of the coexistence equilibrium on $\mathbb{R}^n_{>0}$ directly rules out fold bifurcations at the feasibility boundary.

We therefore seek conditions on $A$ and $B$ under which the coexistence equilibrium of Eq. (1) is globally asymptotically stable on $\mathbb{R}^n_{>0}$.

#### 5.2 Sufficient conditions via the Goh Lyapunov function

To derive such conditions, we employ the classical Goh Lyapunov function [10–12], which is non-negative on $\mathbb{R}^n_{>0}$ and vanishes only at $\mathbf{x}^\star$:

$$V(\mathbf{x}) \;=\; \sum_{i=1}^{n} \left( x_i - x_i^\star - x_i^\star \ln \frac{x_i}{x_i^\star} \right). \tag{22}$$

Along trajectories of Eq. (1), and using $\dot{x}_i = x_i f_i(\mathbf{x})$ together with $f_i(\mathbf{x}^\star) = 0$, the time derivative of $V$ takes the form

$$\dot{V} \;=\; \sum_{i=1}^{n} \frac{x_i - x_i^\star}{x_i}\, \dot{x}_i \;=\; (\mathbf{x}-\mathbf{x}^\star)^\top f(\mathbf{x}) \;=\; (\mathbf{x}-\mathbf{x}^\star)^\top \big(f(\mathbf{x}) - f(\mathbf{x}^\star)\big). \tag{23}$$

Global asymptotic stability on $\mathbb{R}^n_{>0}$ follows if $\dot{V} < 0$ for all $\mathbf{x} \in \mathbb{R}^n_{>0} \setminus \{\mathbf{x}^\star\}$.

***From difference to quadratic form.***

To express $\dot{V}$ as a quadratic form in $(\mathbf{x}-\mathbf{x}^\star)$, we parametrise the straight path from $\mathbf{x}^\star$ to $\mathbf{x}$ as $\mathbf{z}(t) = \mathbf{x}^\star + t(\mathbf{x}-\mathbf{x}^\star)$, so that $\mathbf{z}(0) = \mathbf{x}^\star$, $\mathbf{z}(1) = \mathbf{x}$, and $d\mathbf{z}/dt = \mathbf{x}-\mathbf{x}^\star$. By the fundamental theorem of calculus,

$$f(\mathbf{x}) - f(\mathbf{x}^\star) \;=\; \int_0^1 \frac{d}{dt} f\big(\mathbf{z}(t)\big)\, dt \;=\; \left[\int_0^1 \nabla f\big(\mathbf{x}^\star + t(\mathbf{x}-\mathbf{x}^\star)\big)\, dt\right] (\mathbf{x}-\mathbf{x}^\star). \tag{24}$$

Substituting into Eq. (23) yields the quadratic form

$$\dot{V} \;=\; (\mathbf{x}-\mathbf{x}^\star)^\top \left[\int_0^1 \nabla f\big(\mathbf{x}^\star + t(\mathbf{x}-\mathbf{x}^\star)\big)\, dt\right] (\mathbf{x}-\mathbf{x}^\star). \tag{25}$$

***Evaluating the averaged Jacobian.***

For the GLV+HOI dynamics, and assuming without loss of generality the symmetrisation $B_{ijk} = B_{ikj}$, the per-capita growth rates are $f_i(\mathbf{x}) = r_i + (1-\alpha)\sum_j A_{ij}x_j + \alpha \sum_{j,k} B_{ijk}x_j x_k$, with Jacobian entries

$$\frac{\partial f_i}{\partial x_\ell} \;=\; (1-\alpha)A_{i\ell} + 2\alpha \sum_k B_{i\ell k}\, x_k. \tag{26}$$

Introducing the slice matrices $B_k := (B_{ijk})_{ij}$, this reads in matrix form

$$\nabla f(\mathbf{x}) \;=\; (1-\alpha)\, A \;+\; 2\alpha \sum_{k=1}^{n} x_k\, B_k. \tag{27}$$

Evaluating along the path $\mathbf{z}(t) = \mathbf{x}^\star + t(\mathbf{x}-\mathbf{x}^\star)$ and integrating,

$$\int_0^1 \nabla f\big(\mathbf{z}(t)\big)\, dt \;=\; (1-\alpha)\, A \;+\; \alpha \sum_{k=1}^{n} (x_k + x_k^\star)\, B_k, \tag{28}$$

where we used $\int_0^1 (x_k^\star + t(x_k - x_k^\star))\, dt = (x_k + x_k^\star)/2$. Substituting back into Eq. (25),

$$\dot{V}(\mathbf{x}) \;=\; (\mathbf{x} - \mathbf{x}^\star)^\top \left[ (1-\alpha)\, A \;+\; \alpha \sum_{k=1}^{n} \frac{x_k + x_k^\star}{2}\, B_k \right] (\mathbf{x} - \mathbf{x}^\star). \tag{29}$$

***Sufficient conditions.***

A quadratic form $\mathbf{v}^\top M \mathbf{v}$ is negative for all $\mathbf{v} \neq 0$ iff the symmetric part $S_M := (M + M^\top)/2$ is negative definite. Applying this to the bracketed matrix in Eq. (29), and defining $S_A := (A + A^\top)/2$ and $(S_k)_{ij} := (B_{ijk} + B_{jik})/2$, the map $\mathbf{x} \mapsto \dot{V}(\mathbf{x})$ is strictly negative on $\mathbb{R}^n_{>0} \setminus \{\mathbf{x}^\star\}$ whenever the matrix

$$M(\mathbf{x}) \;=\; (1-\alpha)\, S_A \;+\; \alpha \sum_{k=1}^{n} \frac{x_k + x_k^\star}{2}\, S_k \tag{30}$$

is negative definite for every $\mathbf{x} \in \mathbb{R}^n_{>0}$. Because $(x_k + x_k^\star)/2 > 0$ on the positive orthant, $M(\mathbf{x})$ is a conic (non-negative) combination of $S_A$ and the symmetrised slice matrices $\{S_k\}_{k=1}^n$. A sufficient condition is therefore that each of these matrices be separately negative definite:

$$S_A \;\prec\; 0 \qquad \text{and} \qquad S_k \;\prec\; 0 \quad \text{for every } k = 1, \ldots, n. \tag{31}$$

When condition (31) holds, $\dot{V} < 0$ on $\mathbb{R}^n_{>0} \setminus \{\mathbf{x}^\star\}$ and the coexistence equilibrium $\mathbf{x}^\star$ is globally asymptotically stable on the positive orthant and fold bifurcations are precluded.

The slice-wise condition (31) is strictly stronger than local stability of the coexistence equilibrium. Local stability only constrains the symmetrised community Jacobian evaluated at the single point $\mathbf{x} = \mathbf{x}^\star$, where the slice contributions are weighted by the specific equilibrium abundances $x_k^\star$. In contrast, condition (31) requires the combination in Eq. (30) to be negative definite for *all* $\mathbf{x} \in \mathbb{R}^n_{>0}$. The gap between these two conditions delineates the regime in which multiple equilibria—and hence fold bifurcations—can arise: when local stability holds but the slice-wise condition fails, the interaction structure permits additional equilibria, and feasibility boundaries can contain fold bifurcations of the type documented in the main text.

The same slice-wise condition (31) has been derived in the related context of nonlinear complementarity problems (NCP) [13], where it guarantees uniqueness of the NCP solution via strict monotonicity of the associated mapping $F = -f$ on $\mathbb{R}^n_{>0}$. Since the non-negative equilibria of Eq. (1) are exactly the solutions of the NCP with mapping $F = -f$, the NCP perspective provides an independent route to the same conclusion on uniqueness. The Lyapunov argument given here has the additional advantage of yielding global asymptotic stability—a dynamical statement—rather than uniqueness alone.

### 6 Robustness and generality of fold-mediated collapse

The main-text results rely on a tightly controlled ensemble: the baseline equilibrium is planted at $\mathbf{x}^\star = \mathbf{1}$ for every realization, pairwise and higher-order terms are mixed through a single convex knob $\alpha \in [0, 1]$, the spectral rescaling of Sec. 4 enforces Hurwitz stability at the baseline for all $\alpha$. Each of these controls buys us something: taken together, they isolate the causal effect of HOI strength on feasibility geometry from confounds due to a shifting reference state, a changing interaction magnitude, or an unstable baseline, where ecological dynamics would never settle.

#### 6.1 Quantifying nonlinearity without $\alpha$

Before we can ask whether the main-text pattern survives in other parameterizations or other models, we need a replacement for $\alpha$ that can be evaluated in any system. To this end, this section introduces two model-agnostic measures of nonlinearity—a local one built from derivatives at the coexistence equilibrium, and a nonlocal one built from the geometry of the boundary scan itself. Equipped with these, we then relax the assumptions of the main text in sequence: first the parameterization, keeping the GLV+HOI functional form (Sec. 6.2); then the functional form itself (Sec. 6.3). The same pattern persists once all controls are lifted, illustrating the breadth of our findings.

##### 6.1.1 A local metric.

We define the first metric from a local Taylor expansion of the per-capita growth rate $f$ around the baseline coexistence equilibrium $\mathbf{x}^\star$.

Given any smooth dynamical system $\dot{x}_i = x_i\, f_i(\mathbf{x})$ and a coexistence equilibrium $\mathbf{x}^\star \in \mathbb{R}^n_{>0}$, let

$$J_{ij} \;:=\; \frac{\partial f_i}{\partial x_j}\bigg|_{\mathbf{x}^\star}, \qquad H_{ijk} \;:=\; \frac{\partial^2 f_i}{\partial x_j\, \partial x_k}\bigg|_{\mathbf{x}^\star} \tag{32}$$

denote the Jacobian and Hessian of the per-capita rates at $\mathbf{x}^\star$. Expanding $f$ about $\mathbf{x}^\star$ and collecting, for each species $i$, the first- and second-order contributions evaluated at the equilibrium abundance scale,

$$T_1[i] \;:=\; \sum_j J_{ij}\, x_j^\star, \qquad T_2[i] \;:=\; \tfrac{1}{2} \sum_{j,k} H_{ijk}\, x_j^\star\, x_k^\star, \tag{33}$$

we average their magnitudes across species,

$$P \;:=\; \tfrac{1}{n} \sum_{i=1}^{n} |T_1[i]| \quad \text{(linear magnitude)}, \qquad Q \;:=\; \tfrac{1}{n} \sum_{i=1}^{n} |T_2[i]| \quad \text{(quadratic magnitude)}, \tag{34}$$

and define the Taylor effective nonlinearity as the quadratic share

$$\alpha_{\text{eff}}^{\text{Taylor}} \;:=\; \frac{Q}{P+Q} \;\in\; [0,1]. \tag{35}$$

By construction $\alpha_{\text{eff}}^{\text{Taylor}} = 0$ when $f$ is exactly linear at $\mathbf{x}^\star$ and $\alpha_{\text{eff}}^{\text{Taylor}} \to 1$ as the quadratic response dominates. The metric is a functional of $f$ alone: it is invariant under multiplicative rescalings of $f$ (such as the time rescaling used in Section 6.3 to polynomialise rational systems), independent of how interactions are parameterized, and well-defined for any smooth model.

***Closed form for the main-text parameterization.***

For the main-text construction of Section 4 (Eq. 3) we the constrains so that $\mathbf{x}^\star = \mathbf{1}_n$ is a coexistence equilibrium at $\mathbf{r} = \mathbf{1}_n$ for every $\alpha \in [0,1]$ and for every realization of the random tensors (Eq. (15) followed by the uniform rescaling step). Under this construction, $\alpha_{\text{eff}}^{\text{Taylor}}$ reduces to (See Appendix)

$$\alpha_{\text{eff}}^{\text{Taylor}} = \frac{\alpha}{1+2\alpha}, \qquad \alpha \in [0,1], \tag{36}$$

Note that this nonlinearity proxy is a monotonically increasing function of alpha, confirming that it is indeed a valid measure of HOI strength.

##### 6.1.2 A nonlocal metric

The Taylor-based nonlinearity of is a purely local quantity: $P$ and $Q$ are built from the Jacobian $J$ and Hessian $H$ evaluated *at* the coexistence equilibrium $\mathbf{x}^\star$ (Eqs. (33)–(35)). For systems whose nonlinearity is smooth and global, this pointwise reading of $f$ is informative. For the tipping behaviour we study in the main text, however, it can miss structure that actually drives the collapse. A fold bifurcation is created by the collision of the stable coexistence state with a distinct unstable equilibrium, which typically sits *away* from $\mathbf{x}^\star$ somewhere in the nonnegative orthant; the curvature of $f$ between the two equilibria is what eventually annihilates them as the driver is increased. A system whose per-capita rates look nearly affine at $\mathbf{x}^\star$ can therefore still harbour a fold a finite distance away, and will be assigned a small $\alpha_{\text{eff}}^{\text{Taylor}}$ even when its tipping structure is strongly nonlinear. This gap is not hypothetical: in the closed-form identity Eq. (36) the Taylor metric saturates at $1/3$ as $\alpha \to 1$, because the derivatives at $\mathbf{x}^\star = \mathbf{1}$ cannot see HOI structure that only becomes visible further out in state space.

We therefore introduce a second, nonlocal metric that measures how much of the per-capita rate $f$ over the tipping-relevant region of state space cannot be captured by *any* affine model. Intuitively: if $f$ happens to be affine throughout the region the boundary-scan traverses, the system cannot fold within that region, regardless of how curved $f$ is at isolated points. Conversely, residual variance of $f$ against its best affine approximation provides a direct, coordinate-free measure of the nonlinearity that the fold geometry actually sees.

The boundary-scan of Section 2 sweeps a family of rays outward from $\mathbf{x}^\star$ in environmental space and records, for each ray, the last stable coexistence state $\mathbf{x}_k^{\text{pre}}$ reached before coexistence is lost. We therefore measure non-linearity in this region $\mathcal{H}$.

Let $f : \mathbb{R}^n_{\geq 0} \to \mathbb{R}^n$ denote the per-capita rate at the unperturbed baseline $\mathbf{r} = \mathbf{r}^0$ (no perturbation applied). We draw $M$ samples $\{\mathbf{x}^{(m)}\}_{m=1}^M$ from $\mathcal{H}$ by assigning Dirichlet$(\alpha_{\text{dir}}, \ldots, \alpha_{\text{dir}})$ weights to the hull vertices; the concentration $\alpha_{\text{dir}} = 0.1 < 1$ biases each draw toward a few vertices at a time, so that the sample set as a whole spreads throughout $\mathcal{H}$ rather than clustering near its centroid as a uniform Dirichlet would. The vertices themselves are appended to the sample set so that the extreme preboundary states are always present in the fit; we use 2000 samples.

Stacking the samples into $X \in \mathbb{R}^{n\times(M+|\mathcal{V}|)}$ (where $\mathcal{V}$ is the vertex set) and evaluating the per-capita rates column-wise gives a response matrix $Y \in \mathbb{R}^{n\times(M+|\mathcal{V}|)}$ with $Y_{:,m} = f(\mathbf{x}^{(m)})$. We then fit the best affine model,

$$Y \approx \mathbf{a} + B\,X, \qquad (\hat{\mathbf{a}}, \hat{B}) = \underset{\mathbf{a},B}{\arg\min} \left\| Y - \mathbf{a} - B\,X \right\|_{\text{F}}^2, \tag{37}$$

We define the hull nonlinearity metric as the minimized residual,

$$\alpha_{\text{eff}}^{\text{hull}} := \frac{\sum_{i,m}\left(Y_{im} - \hat{Y}_{im}\right)^2}{\sum_{i,m}\left(Y_{im} - \bar{Y}_i\right)^2} \in [0,1]. \tag{38}$$

By construction $\alpha_{\text{eff}}^{\text{hull}} = 0$ if and only if $f$ is affine on $\mathcal{H}$ (the best linear model captures *all* of its variation), and $\alpha_{\text{eff}}^{\text{hull}} \to 1$ as the residual dominates.

***Validation on the main-text ensemble.***

We now compare our hull-based nonlinearity diagnostic against the main-text ensemble. To this end, we compute $\alpha_{\text{eff}}^{\text{hull}}$ for all the models in the main-text systems, and plot it against $\alpha$ of those systems. The parallel between the two datasets is striking. Across all three $\mu_B$ regimes, the shape of $\alpha_{\text{eff}}^{\text{hull}}(\alpha)$ (Fig. S5a–c, main plots) closely mirrors the shape of fold prevalence as a function of $\alpha$ reported in Fig. 3 of the main text. In the competitive ($\mu_B = -0.1$) and neutral ($\mu_B = 0$) banks, both quantities rise gradually from near zero as $\alpha$ increases, with smaller communities climbing faster than larger ones. In the facilitative bank ($\mu_B = 0.1$), both quantities instead jump almost immediately to a plateau at very small $\alpha$ and remain there, independent of $n$. That $\alpha_{\text{eff}}^{\text{hull}}$—a purely geometric diagnostic computed from the equilibrium manifold without reference to any bifurcation—reproduces the regime-specific qualitative behavior of fold prevalence suggests it captures the geometric feature underlying fold-mediated collapse.

One might then hope to see a tight collapse when plotting fold prevalence directly against $\alpha_{\text{eff}}^{\text{hull}}$ (insets in panels a,b). For the competitive and neutral banks, the relationship is indeed monotonically increasing. In the facilitative bank, the same plot is uninformative: $\alpha_{\text{eff}}^{\text{hull}}$ saturates at $\approx 0.25$–$0.30$ almost as soon as HOIs are turned on, while fold prevalence continues to rise with $\alpha$, so any plot of folds against $\alpha_{\text{eff}}^{\text{hull}}$ would reduce to a nearly vertical band with no resolving power. We therefore replaced the inset in panel c with fold prevalence against $\alpha_{\text{eff}}^{\text{Taylor}}$, which retains variation across the range of $\alpha$ we explore and makes the continued growth of fold prevalence visible.

The need to switch diagnostics between panels is not a deficiency of either measure; rather, it reveals that the two are capturing different regions of nonlinearity and are best read as complementary. The Taylor measure is a *local* quantity: it weighs the pairwise and higher-order contributions to the per-capita growth rate at a single reference equilibrium, and so grows smoothly with $\alpha$ as HOI terms are dialed up in magnitude. The hull measure, by contrast, examines nonlinearity at the further regions of the coexistence domain. As such, the fact that the latter measure does not capture well the fold bifurcation prevalence suggests that the two measures are complementary proxies of the amount of non linearity in the system.

#### 6.2 Robustness to alternative parameterizations

Equipped with these two measures of nonlinearity, we can now test whether fold prevalence is an artifact of the main-text controls rather than of HOIs themselves. To do so, we keep the GLV+HOI functional form but replace the main-text parameterization with the independent constrained-HOI framework of Gibbs et al. [14], which imposes none of our controls: there is no planted $\mathbf{x}^\star = \mathbf{1}$, no convex interpolation knob, and no spectral rescaling; $A$ and $B$ are drawn to *jointly* balance $R$ at a target equilibrium, so neither tensor alone need support feasibility. Because there is no $\alpha$, we quantify nonlinearity post hoc using the two metrics of Sec. 6.1. The main finding

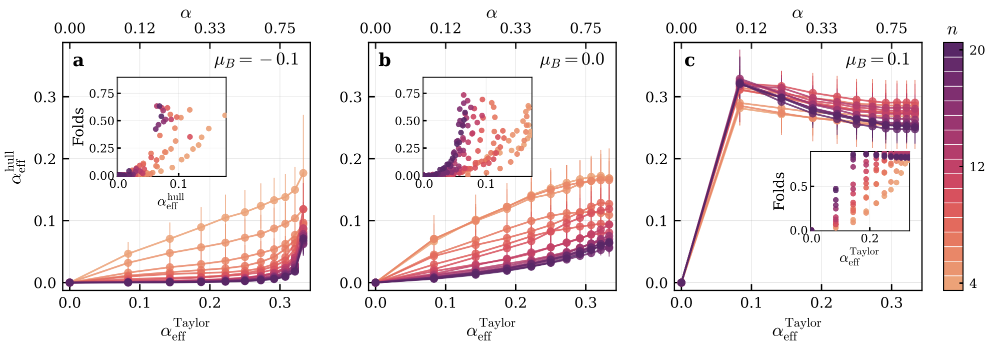


**Fig. S5 Hull nonlinearity vs. Taylor nonlinearity for the three main-text banks.** Mean hull-based effective nonlinearity $\alpha_{\text{eff}}^{\text{hull}}$ (Section 6.1.2, Eq. (38)) plotted against the local Taylor metric $\alpha_{\text{eff}}^{\text{Taylor}}$ (Section 6.1.1, Eq. (35)) for each of the three main-text parameterizations: **(a)** $\mu_B = -0.1$, **(b)** $\mu_B = 0.0$, **(c)** $\mu_B = 0.1$. Each curve corresponds to a fixed diversity $n$ (colors). The top axis (secondary) shows the original mixing parameter $\alpha$ via the closed-form inversion $\alpha = \alpha_{\text{eff}}^{\text{Taylor}}/(1 - 2\alpha_{\text{eff}}^{\text{Taylor}})$ implied by Eq. (36); both $x$-axes therefore index the same grid. Markers and lines show the mean across $\sim 50$ replicates per cell; vertical error bars show $\pm 1$ standard deviation of $\alpha_{\text{eff}}^{\text{hull}}$ across replicates. No horizontal error bars are drawn because $\alpha_{\text{eff}}^{\text{Taylor}}$ is, by the closed-form identity Eq. (36), identical for every replicate of every bank. Insets show the fold prevalence as a function of the nonlocal nonlinearity $\alpha_{\text{eff}}^{\text{hull}}$; capturing a tight correlation for the first two banks, and reflecting the original lack of correlation that alpha had with fold prevalence for the third bank.

persists: fold prevalence is possitively corelated with effective nonlinearity, confirming that fold-mediated coexistence loss is a robust consequence of HOIs rather than an artifact of our controls.

##### 6.2.1 The Gibbs et al. framework

The Gibbs et al. framework starts from the same GLV+HOI dynamics as in the main text, but with sign conventions matching the ecological interpretation directly:

$$\dot{x}_i = x_i\Big(R_i - \sum_j A_{ij}\, x_j - \sum_{j,k} B_{ijk}\, x_j\, x_k\Big), \tag{39}$$

where $R_i > 0$ are intrinsic growth rates, $A_{ij}$ encodes pairwise interactions (with $A_{ij} > 0$ representing competition and the diagonal $A_{ii} > 0$ representing self-regulation), and $B_{ijk}$ represents higher-order interactions. Given a target equilibrium $\mathbf{x}^\star > 0$ and a sampled pairwise matrix $A$, the higher-order tensor $B$ is constrained so that $\mathbf{x}^\star$ satisfies the equilibrium condition:

$$R_i = \sum_j A_{ij}\, x_j^\star + \sum_{j,k} B_{ijk}\, x_j^\star\, x_k^\star, \qquad i = 1, \ldots, n. \tag{40}$$

Crucially, $A$ and $B$ jointly balance $R$ at the target equilibrium—neither component alone need support feasibility. This is fundamentally different from our main-text construction, where $A$ and $B$ are each independently designed so that $\mathbf{x}^\star = \mathbf{1}$ is an equilibrium for all $\alpha$. In addition, this framework has no interpolation parameter $\alpha$; the relative importance of HOIs varies across systems and must be quantified post hoc, using the metrics of Sec. 6.1.

Within this framework, Gibbs et al. identify three qualitatively distinct regimes (Q1, Q2, Q3) in which HOIs commonly produce stable coexistence; each regime

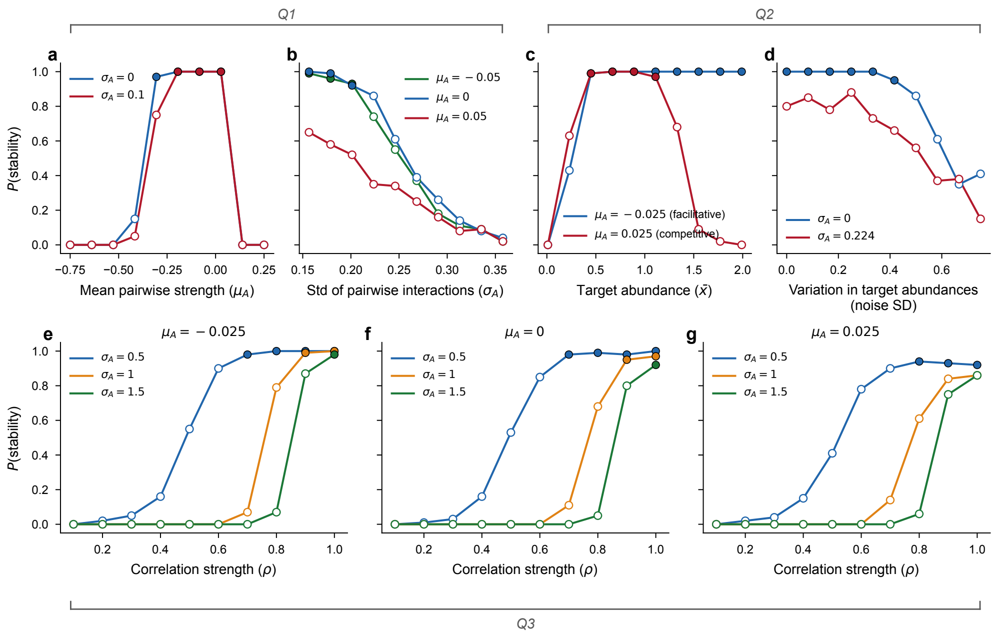


**Fig. S6 Probability of stability across the three Gibbs regimes, replicating the parameter sweeps of Gibbs et al. with $S = 20$ species.** (a, b) Regime Q1: P(stability) as a function of mean pairwise interaction strength $\mu_A$ (a) and standard deviation $\sigma_A$ (b), with HOI coefficients constrained via Dirichlet sampling. (c, d) Regime Q2: P(stability) as a function of target abundance $\bar{x}$ (c) and noise in target abundances (d). (e–g) Regime Q3: P(stability) as a function of anti-correlation strength $p$ for three values of $\mu_A$, with curves colored by $\sigma_A \in \{0.5, 1.0, 1.5\}$. Filled circles indicate parameterizations with $P(\text{stability}) > 0.9$; open circles denote those below the threshold.

involves different pairwise statistics, target abundances, and stabilization mechanisms. We generated systems in all three, providing natural variation in interaction structure while staying within parameter regions where stability is common. Full regime specifications are given in the Appendix.

##### 6.2.2 System generation and acceptance

For each parameter combination we generated candidate systems at community size $S = 20$ by sampling pairwise interactions $A$ and computing the constrained HOI tensor $B$ as described above, using growth rates $R_i = 1$ and target abundances $\mathbf{x}^\star$ determined by the regime. We restricted generation to parameter combinations where the probability of stability (estimated from 100 replicates per combination) exceeds 0.9. This threshold ensures a high acceptance rate during rejection sampling: because each candidate system is drawn from the prescribed distributions over $A$ and $B$ and then accepted or discarded based on a Jacobian eigenvalue check, a low acceptance rate would mean that surviving systems are a heavily filtered subset, distorting the statistical structure of the interactions. By operating in the high-acceptance region of parameter space, we ensure that accepted systems are typical draws from the Gibbs constrained distributions rather than rare outliers that happen to be stable.

Figure S6 shows the probability of stability across the three regimes, replicating the parameter sweeps of Gibbs et al. Filled dots indicate the parameterizations meeting the $P(\text{stability}) > 0.9$ threshold used to populate the system bank.

###### 6.2.3 Boundary detection and results

Given the bank of stable systems produced by the procedure above, we tested each one for fold-mediated coexistence loss. For each system we set up the polynomial equilibrium system

$$(r_i^0 + \delta\, u_{ik}) - \sum_j A_{ij}\, x_j - \sum_{j,k} B_{ijk}\, x_j\, x_k = 0, \qquad i = 1, \ldots, n, \tag{41}$$

where $k$ index the perturbation direction, and applied the boundary-detection algorithm described in Sec. 2. Figure S7 shows the results. Because we have dropped all controls—working with an orthogonal parameterization, exploring parameter combinations with different interaction statistics than in the main text, and computing nonlinearity post hoc—the results are expectedly noisier than our controlled simulations. Nevertheless, the central pattern is clear: for some parameterizations, folds become very common, and these sit at high $\alpha_{\text{eff}}$. This confirms the generality of our finding that fold prevalence increases with the nonlinearity of the system.

##### 6.3 Robustness to alternative functional forms

This step requires a preliminary observation. The higher-order tensor $B_{ijk}$ of the main text is a phenomenological object: it captures the net effect of one species on the interaction between two others, but does not specify the mechanism producing that effect. What experiments and ecological models routinely document instead are nonlinear functional responses—saturating uptake, handling-time constraints, density-dependent interference—that make per-capita interaction effects depend on community state [15, 16]. As such, in this section we now loosen the functional form itself and ask whether fold-mediated collapse remains common in published multi-species models that were not built around a three-way interaction tensor—models with saturating mutualistic responses, allometric trophic networks, ecosystem-engineering feedbacks, Allee effects, and Holling-type predation.

We start by showing that the two descriptions are formally equivalent: any pairwise model whose per-capita growth rates are rational or smooth functions of abundances can be recast, exactly or locally, as a GLV model with effective higher-order interactions. Consider a general ecological model:

$$\dot{x}_i = x_i\, g_i(\mathbf{x}), \tag{42}$$

where each $g_i$ can be written as a ratio of polynomials. Let $S(\mathbf{x}) > 0$ denote the product of all denominators appearing across every $g_i$ (which is strictly positive for positive abundances). We rescale time globally by

$$d\tau = S(\mathbf{x})\, dt.$$

Since $S(\mathbf{x}) > 0$ for all $\mathbf{x} > 0$, $\tau$ is strictly increasing along trajectories, and the chain rule gives

$$\frac{dx_i}{d\tau} = x_i\, S(\mathbf{x})\, g_i(\mathbf{x}), \tag{43}$$

where the product $S(\mathbf{x})\, g_i(\mathbf{x})$ is now a polynomial in $\mathbf{x}$ for every $i$, since all denominators have been cleared. Crucially, because $S(\mathbf{x}) > 0$ for all $\mathbf{x} > 0$, the equilibria of the rescaled system (43) coincide exactly with those of the original dynamics (42): multiplying every right-hand side by the same strictly positive factor $S$ does not

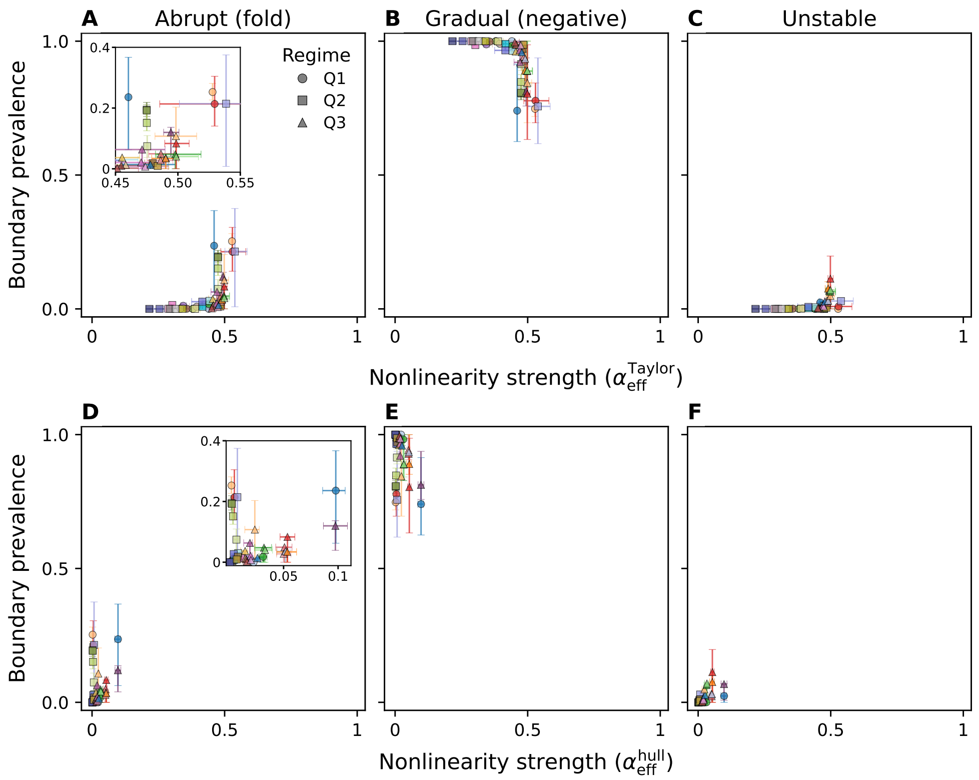


**Fig. S7 Boundary-type prevalence vs. effective nonlinearity strength in the Gibbs bank ($n$ = 20).** Each point aggregates one parameterization across its replicates (47 parameterizations, 4612 stable systems, 128 random parameter rays per system). Marker position gives the means of the effective nonlinearity $\alpha_{\text{eff}}$ on the $x$-axis and the fraction of rays terminating in the indicated boundary type on the $y$-axis; error bars span the 25th–75th percentile across replicates. Marker shape encodes the parameter regime, color encodes the parameterization id (filled circles in Fig. S6. **A, D:** fraction of rays ending at a fold (abrupt loss of coexistence). **B, E:** fraction ending at a negative-crossing boundary (gradual loss). **C, F:** fraction ending at a loss-of-stability boundary. **Top row (A–C):** $x$-axis is the Taylor-expansion estimate $\alpha_{\text{eff}}^{\text{Taylor}}$. **Bottom row (D–F):** $x$-axis is the convex-hull estimate $\alpha_{\text{eff}}^{\text{hull}}$.

create or destroy any zeros. Therefore, the bifurcation structure—including the existence and location of fold bifurcations at the feasibility boundary—is identical in both formulations.

This transformation shows that any model with saturating nonlinearities is, from the standpoint of equilibrium analysis, a GLV model with higher-order interactions—potentially of very high order.

An alternative route to effective higher-order interactions does not require the per-capita growth rates to be rational. For any smooth nonlinear model of the form (42), we can Taylor-expand each per-capita growth rate $g_i(\mathbf{x})$ around a reference equilibrium $\mathbf{x}^\star$:

$$g_i(\mathbf{x}) = g_i(\mathbf{x}^\star) + \sum_j \frac{\partial g_i}{\partial x_j}\bigg|_{\mathbf{x}^\star} (x_j - x_j^\star) + \frac{1}{2}\sum_{j,k} \frac{\partial^2 g_i}{\partial x_j \partial x_k}\bigg|_{\mathbf{x}^\star} (x_j - x_j^\star)(x_k - x_k^\star) + \cdots \quad (44)$$

The zeroth-order term gives the intrinsic growth rate at equilibrium, the first-order derivatives yield the standard pairwise interaction coefficients, and the second-order derivatives define effective three-way higher-order interactions $B^{\text{eff}}_{ijk} = \frac{1}{2}\,\partial^2 g_i/\partial x_j \partial x_k|_{\mathbf{x}^\star}$. Higher-order terms in the expansion correspond to interactions of progressively higher order. Truncating at second order recovers the GLV+HOI model in Eq. (1), with effective interaction tensors $A$ and $B$ determined by the local curvature of the nonlinear functional responses. This approach is more general than the time-rescaling method above—it applies to any sufficiently smooth model, not only those with rational nonlinearities—but it is inherently local: the effective HOI coefficients depend on the reference equilibrium, and the approximation degrades far from it. The two approaches are thus complementary: the time-rescaling provides an exact, global polynomial reformulation when the nonlinearities are rational, while the Taylor expansion provides a local HOI approximation for arbitrary smooth nonlinearities.

#### 6.4 Published model specifications and parameterizations

Based on the connection between nonlinearities and effective HOIs established above, we apply our boundary-scan algorithm to several published pairwise models with nonlinear functional responses. For each model the procedure is the same:

1. **Sample a feasible, stable equilibrium.** We draw parameters from the distributions specified by the original study and numerically locate a coexistence equilibrium $\mathbf{x}^\star > 0$ that is locally stable through numerical integration, discarding replicates that fail to converge.
2. **Polynomialise.** Because our homotopy-continuation (HC) scanner requires a polynomial system, we clear denominators in every per-capita growth equation to obtain an equivalent polynomial system $\mathbf{G}(\mathbf{x}; \boldsymbol{\delta r}) = \mathbf{0}$, where $\boldsymbol{\delta r} = \delta\,\mathbf{u}$ is the parameter perturbation and $\mathbf{u}$ is a unit direction drawn from the appropriate subspace. Clearing denominators can in principle introduce spurious solutions that are absent in the original system; however, because we always track an equilibrium branch seeded at a feasible equilibrium of the original dynamics, those extra solutions remain outside our computational scope.
3. **Perturb and track.** Starting from $\mathbf{x}^\star$, we use coefficient-parameter homotopy continuation to track the coexistence branch as $\delta$ increases, classifying the first boundary encountered as gradual, abrupt, or unstable using the algorithm described in Sec. 2.

Unlike the canonical GLV-HOI models studied in the main text, where a mixing parameter $\alpha$ interpolates between pairwise and higher-order regimes, these published models possess emergent nonlinearity that cannot be tuned up or down. We therefore treat each model as a single data point at a fixed value of the two measures of effective higher-order strength $\alpha_{\text{eff}}$ (defined above) rather than scanning over an $\alpha$-grid. The figure with results is presented in Fig. S8

The contrast between the two panels of Fig. S8 is itself diagnostic: the Taylor metric, being local, compresses models whose nonlinearity lives away from $\mathbf{x}^\star$ – notably the facilitation-dominated Aguadé-Gorgorío systems, which saturate near the upper range of $\alpha^{\text{Taylor}}_{\text{eff}}$ despite exhibiting fold fractions close to unity – whereas the hull metric, which samples $f$ across the pre-boundary region, separates these systems along the $x$-axis in accordance with their fold prevalence. The two panels therefore confirm empirically what the construction of the metrics predicts: each diagnostic resolves a different regime of nonlinearity, and their joint use is what makes the cross-model comparison interpretable.

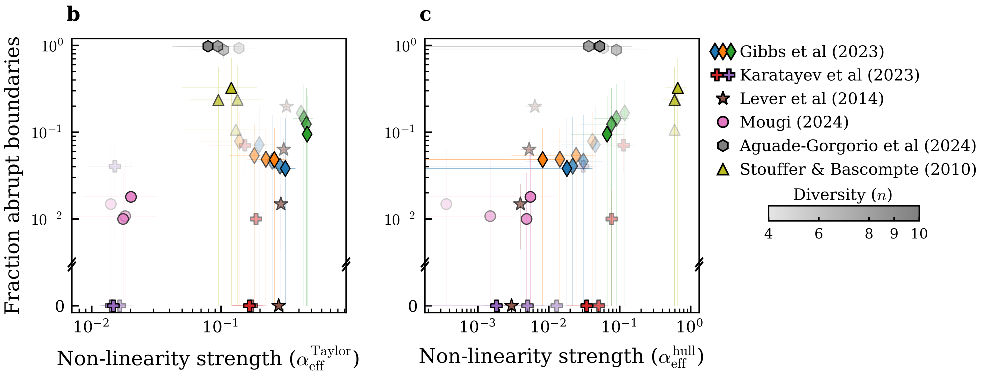


**Fig. S8 Fraction of abrupt (fold) boundaries versus two alternative non-linearity metrics.** Each marker is a bank of models grouped by family and diversity $n$; vertical and horizontal bars show the standard deviation of the fold fraction and of the non-linearity metric within the group, respectively. (**b**) Non-linearity strength measured by the Taylor-expansion unified metric $\alpha_{\text{eff}}^{\text{Taylor}}$. (**c**) Non-linearity strength measured by $\alpha_{\text{eff}}^{\text{hull}}$, the $L^2$ fraction of the nominal vector field contributed by higher-order terms, integrated over the convex hull of the stable equilibrium and the pre-boundary states. Marker shape and colour identify the model family (legend, right); marker opacity encodes species richness $n$ (colourbar, right). Both panels share the same $y$-axis.

The remainder of the section provides the full dynamical equations, parameterizations, and perturbation protocols used for each of the five published models analysed in this study.

##### 6.4.1 Plant-pollinator network (Lever et al, 2014)

***Original model.***

The community comprises $S_P$ plant species and $S_A$ pollinator (animal) species [17], indexed $i = 1, \ldots, S_P$ and $k = 1, \ldots, S_A$, respectively. Species ordering in the state vector is $\mathbf{x} = (P_1, \ldots, P_{S_P}, A_1, \ldots, A_{S_A})^\top$, with $n = S_P + S_A$.

Plant dynamics:

$$\frac{dP_i}{dt} = P_i \left[ r_i^P + \frac{\sum_{k=1}^{S_A} \gamma_{ik}^P A_k}{1 + h_i^P \sum_{k=1}^{S_A} \gamma_{ik}^P A_k} - \sum_{j=1}^{S_P} c_{ij}^P P_j \right] + \mu_P \,, \tag{45}$$

Pollinator dynamics:

$$\frac{dA_k}{dt} = A_k \left[ r_k^A - d_A + \frac{\sum_{i=1}^{S_P} \gamma_{ki}^A P_i}{1 + h_k^A \sum_{i=1}^{S_P} \gamma_{ki}^A P_i} - \sum_{l=1}^{S_A} c_{kl}^A A_l \right] + \mu_A \,. \tag{46}$$

The bipartite interaction topology is fully connected ($\gamma_{ik}^P, \gamma_{ki}^A > 0$ for all plant–pollinator pairs). Mutualistic benefit coefficients are normalised by partner degree via the trade-off parameter $\theta$:

$$\gamma_{ik}^P = \frac{\gamma_{ik}^{P,0}}{d_i^\theta} \,, \qquad \gamma_{ki}^A = \frac{\gamma_{ki}^{A,0}}{d_k^\theta} \,, \tag{47}$$

where $d_i = \sum_k \mathbf{1}[\gamma_{ik}^P > 0]$ is the degree of plant $i$ (and analogously for pollinators), and $\gamma_{ik}^{P,0}, \gamma_{ki}^{A,0}$ are baseline strengths. For this model, we resolved the bifurcation structure explicitly.

***Bifurcation diagram construction (Fig. 4A).***

To resolve the full bifurcation structure of the Lever et al. model explicitly, we construct the diagram in Fig.4A as follows. We set up a parameter homotopy in `HomotopyContinuation.jl` that varies pollinator mortality $d_A$ from 0 (the baseline) to a target value, holding all other parameters fixed. Starting from the numerically obtained coexistence equilibrium at $d_A = 0$, we track the stable coexistence with homotopy continuation using $d_A$ as perturbation parameter. The tracker terminates when the Jacobian $\frac{\mathrm{d}H}{\mathrm{d}\mathbf{x}}$ becomes singular, indicating a fold bifurcation at a critical mortality $d_A^c$. To locate the unstable branch that meets the stable branch at this fold, we evaluate the system at a parameter value slightly before the fold ($d_A \lesssim d_A^c$), compute the near-singular direction of the Jacobian via SVD, and use it to seed a Newton search for a second, distinct equilibrium. Once found, this unstable equilibrium is continued backward (toward $d_A = 0$) using the same parameter homotopy, yielding the unstable branch. To confirm that the stable algebraic branch correspond to realized dynamics, we overlay numerical integration of the original ODE system (45)–(46). A forward sweep integrates the system to steady state at each of a sequence of $d_A$ values (using the previous endpoint as the initial condition), tracing the stable branch until the fold, after which the system collapses to a zero-abundance attractor. A backward sweep then integrates from the collapsed state as $d_A$ is reduced, revealing hysteresis: recovery occurs at a $d_A$ value corresponding to the terminus of the unstable branch, well below $d_A^c$. The combined algebraic branches (stable and unstable, shown as solid and dashed lines) together with the forward and backward integration endpoints (shown as triangles) constitute the bifurcation diagram in Fig. 4A.

***Parameterization.***

All parameters are drawn independently for each replicate from the distributions listed in Table 6.4.1. Only replicates converging to a strictly positive equilibrium ($x_i^* > 10^{-2}$ for all $i$) are retained.

| Parameter | Description | Distribution | Value / Range |
|---|---|---|---|
| $r_i^P$, $r_k^A$ | Intrinsic growth rates | Uniform | [0.05, 0.35] |
| $h_i^P$, $h_k^A$ | Handling times (saturating response) | Uniform | [0.15, 0.30] |
| $c_{ij}^P$ $(i \neq j)$ | Interspecific competition (plants) | Uniform | [0.01, 0.05] |
| $c_{ii}^P$ | Intraspecific competition (plants) | Uniform | [0.80, 1.10] |
| $c_{kl}^A$ $(k \neq l)$ | Interspecific competition (pollinators) | Uniform | [0.01, 0.05] |
| $c_{kk}^A$ | Intraspecific competition (pollinators) | Uniform | [0.80, 1.10] |
| $\gamma_{ik}^{P,0}$ | Baseline mutualistic benefit (plant $\leftarrow$ pollinator) | Uniform | [0.80, 1.20] |
| $\gamma_{ki}^{A,0}$ | Baseline mutualistic benefit (pollinator $\leftarrow$ plant) | Uniform | [0.80, 1.20] |
| $\theta$ | Degree trade-off exponent | Fixed | 0.5 |
| $\mu_P$, $\mu_A$ | Immigration | Fixed | 0 |
| $d_A$ | Additional pollinator mortality | Fixed | 0 |

***Perturbation protocol.***

Perturbation directions $\mathbf{u} \in \mathbb{R}^n$ are constructed with *zero entries for plant species* and unit-normalised random Gaussian entries in the pollinator subspace: $\mathbf{u}$ =

$(0,\ldots,0,\, u_{S_P+1},\ldots,u_n)^\top$ with $(u_{S_P+1},\ldots,u_n) \sim \mathcal{N}(0, I_{S_A})$ normalised to unit length. This reflects the ecological scenario in which pollinator mortality is the driver of community collapse, consistent with the original study.

***Replicates.***

50 parameter replicates per species richness, 128 perturbation directions per replicate. Community sizes: $n \in \{4, 6, 8, 10\}$ (with $S_P = S_A = n/2$).

##### 6.4.2 Resource-consumer food web with density-mediated feedbacks (Karatayev et al, 2023).

***Original model.***

The community consists of $n_R$ resource species $N_i$ $(i = 1,\ldots,n_R)$ and $n_C$ consumer species $C_k$ $(k = 1,\ldots,n_C)$ [18], with $n = n_R + n_C$ and state vector $\mathbf{x} = (N_1,\ldots,N_{n_R}, C_1,\ldots,C_{n_C})^\top$.

Two feedback modes are considered:

**FMI (Feeding-Mediating Interactions).** Resource species become less edible at high density, creating a density-dependent feeding interaction.

Resource dynamics:

$$\frac{dN_i}{dt} = N_i\left[r_i\left(1 - \frac{(1-a)\,N_i + a\sum_j N_j}{K_i}\right) - \sum_k \sigma_{ik}\left(1 - \frac{f_i\,N_i}{K_i}\right)\delta_k\,C_k\right], \tag{48}$$

Consumer dynamics:

$$\frac{dC_k}{dt} = C_k\left[\delta_k\,b_k \sum_i \sigma_{ik}\left(1 - \frac{f_i\,N_i}{K_i}\right)N_i - m\right] - \beta_k\,C_k^2\,. \tag{49}$$

**RMI (Recruitment-Mediating Interactions).** Resource density modulates consumer recruitment efficiency rather than edibility.

Resource dynamics:

$$\frac{dN_i}{dt} = N_i\left[r_i\left(1 - \frac{(1-a)\,N_i + a\sum_j N_j}{K_i}\right) - \sum_k \sigma_{ik}\,\delta_k\,C_k\right], \tag{50}$$

Consumer dynamics:

$$\frac{dC_k}{dt} = C_k\left[\delta_k\,b_k\left(\sum_i \sigma_{ik}\,N_i\right)\left(1 - \sum_i \omega_{ik}\frac{f_i\,N_i}{K_i}\right) - m\right] - \beta_k\,C_k^2\,. \tag{51}$$

***Parameterization.***

Parameters are sampled following the defaults of [18] (Table 6.4.2). All $\xi$ draws are independent Uniform$(-1, 1)$. The consumer preference matrix $\sigma_{ik}$ is drawn column-wise from a log-normal distribution with $\sigma_{\log} = 0.5$ and then column-normalised. The recruitment weight matrix $\omega_{ik}$ (RMI mode) is drawn row-wise from a Gamma(1) distribution and row-normalised. Resources and consumers are split evenly: $n_R = \lfloor n/2 \rfloor$, $n_C = n - n_R$.

| Parameter | Description | Baseline | Sampling |
|---|---|---|---|
| $K_i$ | Resource carrying capacity | $\bar{K} = 1.35$ | $\bar{K}\,(1 + 0.15\,\xi_i)$ |
| $r_i$ | Resource intrinsic growth rate | $\bar{r} = 1.0$ | $\bar{r}\,(1 + 0.15\,\xi_i)$ |
| $\delta_k$ | Consumer grazing rate | $\bar{\delta} = 1.1$ | $\bar{\delta}\,(1 + 0.15\,\xi_k)$ |
| $b_k$ | Consumer conversion efficiency | $\bar{b} = 1.0$ | $\bar{b}\,(1 + 0.15\,\xi_k)$ |
| $\beta_k$ | Consumer density dependence | $\bar{\beta} = 0.15$ | $\bar{\beta}\,(1 + 0.15\,\xi_k)$ |
| $f_i$ | Edibility/recruitment feedback | $\bar{f} = 0.8$ | $\bar{f}\,(1 + 0.125\,\xi_i)$ |
| $m$ | Baseline consumer mortality | 0.025 | Fixed |
| $a$ | Interspecific resource competition | 0.025 | Fixed |
| $\sigma_{ik}$ | Diet specialisation matrix | — | LogNormal, column-normalised |
| $\omega_{ik}$ | Recruitment weight matrix (RMI) | — | Gamma(1), row-normalised |

***Perturbation protocol.***

Perturbation directions have zero entries for resource species and unit-normalised Gaussian entries in the consumer subspace: $\mathbf{u} = (0, \ldots, 0, u_{n_R+1}, \ldots, u_n)^\top$, reflecting consumer mortality as the perturbation parameter.

***Replicates.***

50 replicates per species richness, 128 directions per replicate, for each of FMI and RMI feedback modes. Community sizes: $n \in \{4, 6, 8, 10\}$ (i.e. $n_R = n_C \in \{2, 3, 4, 5\}$).

##### 6.4.3 Ecosystem-engineering food web (Mougi, 2024)

***Original model.***

The model [19] describes an $n$-species food web with directed trophic links, where a subset of "engineer" species modify the environment and thereby alter the growth rates and interaction strengths experienced by "receiver" species. These engineering effects create effective higher-order interactions.

Species dynamics:

$$\frac{dx_i}{dt} = x_i \left[ r_i\,\mathcal{B}_i(\mathbf{x}) - s_i\,x_i + \mathcal{G}_i(\mathbf{x})\,\frac{\sum_j e_{ij}\,a_{ij}\,x_j}{1 + \sum_j h_{ij}\,a_{ij}\,x_j} - \sum_{\text{pred}\,p} \frac{\mathcal{G}_p(\mathbf{x})\,a_{pi}\,x_p}{1 + \sum_j h_{pj}\,a_{pj}\,x_j} \right], \tag{52}$$

where $\mathcal{B}_i$ and $\mathcal{G}_i$ are engineering modifications of the growth rate and trophic interaction strength, respectively. The engineering factors take the form

$$\mathcal{B}_i(\mathbf{x}) = 1 + \sum_{\text{eng}\,e} (\beta_{ei} - 1)\,\frac{x_e}{x_e + E_e^0}\,, \qquad \mathcal{G}_i(\mathbf{x}) = 1 + \sum_{\text{eng}\,e} (\gamma_{ei} - 1)\,\frac{x_e}{x_e + E_e^0}\,, \tag{53}$$

where the sums run over engineer species $e$, and $\beta_{ei}, \gamma_{ei}$ are the engineering coefficients ($> 1$: facilitation, $< 1$: inhibition), and $E_e^0 > 0$ is a half-saturation constant for the engineering effect. Non-receiver species have $\mathcal{B}_i = \mathcal{G}_i = 1$.

***Parameterization.***

Parameters are sampled as listed in Table 6.4.3. The trophic network is a random directed graph with connectance 1.0 (fully connected): for each unordered pair $(i, j)$ with $i < j$, a directed trophic link is created with random orientation.

The numbers of engineer and receiver species are chosen so that the "engineering dominance" $(n_{\text{eng}}/n) \times (n_{\text{rec}}/n)$ lies in $[0.08, 0.20]$. Engineering coefficients for each engineer–receiver pair: with probability $q_r \in [0.1, 0.3]$, $\beta_{ei} \sim \text{Uniform}(0, 1)$ (inhibition), otherwise $\text{Uniform}(1, 2)$ (facilitation); with probability $q_a \in [0.7, 0.9]$, $\gamma_{ei} \sim \text{Uniform}(0, 1)$, otherwise $\text{Uniform}(1, 2)$.

| Parameter | Description | Distribution | Value / Range |
|---|---|---|---|
| $r_i$ | Baseline intrinsic growth rate | Uniform | [0.2, 1.0] |
| $s_i$ | Self-regulation coefficient | Fixed | 1.0 |
| $a_{ij}$ | Attack rate (if trophic link exists) | Uniform | [0.0, 0.01] |
| $e_{ij}$ | Assimilation efficiency | Uniform | [0.1, 0.25] |
| $h_{ij}$ | Handling time (Type II response) | Uniform | [1.0, 10.0] |
| $E_e^0$ | Engineering half-saturation | Uniform | [0.0, 0.1] |
| $\beta_{ei}$ | Engineering effect on growth rate | See text | $[0, 2]$ |
| $\gamma_{ei}$ | Engineering effect on trophic strength | See text | $[0, 2]$ |

***Perturbation protocol.***

Perturbation directions are full-dimensional: $\mathbf{u} \sim \mathcal{N}(\mathbf{0}, I_n)$, normalised to unit length. The perturbation $\boldsymbol{\delta r} = \delta\,\mathbf{u}$ is added to the baseline growth rates $r_i$.

***Replicates.***

50 replicates per species richness, 128 directions per replicate. Community sizes: $n \in \{4, 6, 8, 9\}$.

##### 6.4.4 Facilitation-competition community with Allee effects (Aguadé-Gorgorió et al. 2024)

***Original model.***

The model [20] describes $n$ species with saturating facilitation (generating Allee-like effects) and linear competition:

$$\frac{dx_i}{dt} = x_i \left[ \sum_{j=1}^{n} \frac{A_{ij}\, x_j}{\gamma_j + x_j} - d_i - \sum_{j=1}^{n} B_{ij}\, x_j \right], \tag{54}$$

where $A_{ij} \geq 0$ is a facilitation matrix (diagonal: Allee-type self-facilitation; off-diagonal: mutualism), $B_{ij} \geq 0$ is a competition matrix (diagonal: self-regulation; off-diagonal: interspecific competition), $\gamma_j > 0$ are half-saturation constants, and $d_i > 0$ are per-species death rates (perturbation parameters in the HC scan). The saturating facilitation generates effective higher-order interactions when expanded around equilibrium.

***Parameterization.***

Parameters follow the protocol of [20], using log-normal distributions (Table 6.4.4). All matrices are dense (no sparsity masking). The interspecific means $\bar{A}$ and $\bar{B}$, as well as the heterogeneity $\sigma_{\text{inter}}$, are drawn once per replicate; individual $A_{ij}$, $B_{ij}$ are then drawn from the corresponding log-normal.

| Parameter | Description | Distribution | Mean / Range |
|---|---|---|---|
| *Intraspecific (sampled independently per species)* | | | |
| $\gamma_i$ | Half-saturation constant | $\mathrm{LogNormal}\big(\ln(\bar{\gamma}),\ \ln(1.1)\big)$ | $\bar{\gamma} = 1.0$ |
| $d_i$ | Death rate | $\mathrm{LogNormal}\big(\ln(\bar{d}),\ \ln(1.1)\big)$ | $\bar{d} = 0.1$ |
| $A_{ii}$ | Self-facilitation (Allee) | $\mathrm{LogNormal}\big(\ln(\bar{A}_{\mathrm{diag}}),\ \ln(1.1)\big)$ | $\bar{A}_{\mathrm{diag}} = 0.5$ |
| $B_{ii}$ | Self-regulation | $\mathrm{LogNormal}\big(\ln(\bar{B}_{\mathrm{diag}}),\ \ln(1.1)\big)$ | $\bar{B}_{\mathrm{diag}} = 0.1$ |
| *Interspecific (means drawn once per replicate)* | | | |
| $A_{ij}$ $(i \neq j)$ | Mutualistic facilitation | $\mathrm{LogNormal}\big(\ln(\bar{A}),\ \ln(\sigma_{\mathrm{inter}})\big)$ | $\bar{A} \sim \mathrm{Uniform}(0^+, 0.5)$ |
| $B_{ij}$ $(i \neq j)$ | Interspecific competition | $\mathrm{LogNormal}\big(\ln(\bar{B}),\ \ln(\sigma_{\mathrm{inter}})\big)$ | $\bar{B} \sim \mathrm{Uniform}(0^+, 0.14)$ |
| $\sigma_{\mathrm{inter}}$ | Interaction heterogeneity | $\mathrm{Uniform}(1.00001,\ 3.0001)$ | — |

***Perturbation protocol.***

Perturbation directions are full-dimensional: $\mathbf{u} \sim \mathcal{N}(\mathbf{0}, I_n)$, normalised to unit length. The perturbation is added to the death rates: $d_i \to d_i + \delta\, u_i$.

***Replicates.***

50 replicates per species richness, 128 directions per replicate. Community sizes: $n \in \{4, 6, 8, 10\}$.

##### 6.4.5 Allometric trophic network (Stouffer & Bascompte, 2010)

***Original model.***

The model [21] uses a niche-model food web [22] with allometrically scaled metabolic rates and Holling Type II functional responses. The state variable $B_i$ denotes the biomass of species $i$.

Basal (producer) species:

$$\frac{dB_i}{dt} = B_i\left[1 - \frac{\sum_{\mathrm{basal}\,j} B_j}{K}\right] - \sum_{\mathrm{pred}\,p} \frac{x_p\, y_p\, w_{pi}\, B_p\, B_i}{e_{pi}\Big(B_0 + \sum_{\mathrm{prey}\,j'} w_{pj'}\, B_{j'}\Big)}, \tag{55}$$

Consumer species:

$$\frac{dB_i}{dt} = B_i\left[-x_i + \frac{x_i\, y_i \sum_{\mathrm{prey}\,j} w_{ij}\, B_j}{B_0 + \sum_{\mathrm{prey}\,j} w_{ij}\, B_j}\right] - \sum_{\mathrm{pred}\,p} \frac{x_p\, y_p\, w_{pi}\, B_p\, B_i}{e_{pi}\Big(B_0 + \sum_{\mathrm{prey}\,j'} w_{pj'}\, B_{j'}\Big)}, \tag{56}$$

where $x_i = (a_x/a_r)\,(M_i/M_b)^{-0.25}$ is the mass-specific metabolic rate of consumer $i$ (allometrically scaled), $y_i = a_y/a_x$ is the maximum ingestion rate relative to metabolic rate, $w_{ij}$ is the attack rate of consumer $i$ on prey $j$, $e_{ij}$ is the assimilation efficiency, $M_i$ is the body mass of species $i$, $K$ is the shared basal carrying capacity, and $B_0$ is the half-saturation constant of the functional response.

***Parameterization.***

Parameter values are listed in Table 6.4.5. The food web topology is generated using the niche model [22]

| Parameter | Description | Distribution | Value |
|---|---|---|---|
| $K$ | Basal carrying capacity | Fixed | 1.0 |
| $B_0$ | Functional-response half-saturation | Fixed | 0.5 |
| $M_b$ | Basal body mass | Fixed | 1.0 |
| $a_r$ | Basal mass-specific metabolic rate | Fixed | 1.0 |
| $a_x$ | Consumer mass-specific metabolic rate | Fixed | 0.314 |
| $a_y$ | Maximum ingestion rate constant | Fixed | $8\,a_x = 2.512$ |
| $e_{ij}$ | Assimilation efficiency | Fixed | 0.85 |
| $w_{ij}$ | Attack rate per trophic link | LogNormal$(-3.0,\ 1.5)$ | — |
| $M_i$ | Consumer body mass | MCMC (see text) | — |

***Perturbation protocol.***

Perturbation directions are full-dimensional: $\mathbf{u} \sim \mathcal{N}(\mathbf{0}, I_n)$, normalised to unit length. For basal species, the perturbation shifts the intrinsic growth rate: $1 + \delta r_i$ replaces 1 in Eq. (55). For consumers, the perturbation shifts the metabolic rate: $-x_i + \delta r_i$ replaces $-x_i$ in Eq. (56).

***Replicates.***

50 replicates per species richness, 128 directions per replicate. Community sizes: $n \in \{4, 6, 8, 10\}$.

### Appendix A  Mean matrices of $\boldsymbol{S_A}$ and $\boldsymbol{S_{\hat{B}}}$

We compute the mean matrices for both the pairwise symmetric matrix $S_A = \frac{1}{2}(A + A^\top)$ and the symmetric HOI aggregation matrix $(S_{\hat{B}})_{ij} = \frac{1}{4}\sum_k (B_{ijk} + B_{ikj} + B_{jik} + B_{jki})$, working before the diagonal rescaling in Eq. (17). In both cases, the zero-row-sum (resp. zero-slice-sum) constraint creates a deterministic coupling between off-diagonal and diagonal entries that shifts the spectrum away from the origin whenever the interaction mean is nonzero.

#### Pairwise sector: $\mathbb{E}[\boldsymbol{S_A}]$

Recall that $\tilde{A}_{ij} \sim \mathcal{N}(\mu_A, n^{-1})$ for $i \neq j$, with the zero-row-sum constraint $\tilde{A}_{ii} = -\sum_{j\neq i} \tilde{A}_{ij}$.

***Off-diagonal* $(i \neq j)$*.***
The entries $\tilde{A}_{ij}$ and $\tilde{A}_{ji}$ are independent unconstrained draws, so

$$\mathbb{E}[(S_A)_{ij}] = \tfrac{1}{2}\,\mathbb{E}[\tilde{A}_{ij} + \tilde{A}_{ji}] = \mu_A.$$

***Diagonal* $(i = j)$*.***
By the zero-sum constraint,

$$\mathbb{E}[(S_A)_{ii}] = \mathbb{E}[\tilde{A}_{ii}] = -(n-1)\mu_A.$$

Combining:

$$\mathbb{E}[S_A] = \mu_A\, \mathbf{1}\mathbf{1}^\top - n\mu_A\, I,$$

which factors as

$$\mathbb{E}[S_A] = -n\mu_A \left( I - \frac{\mathbf{1}\mathbf{1}^\top}{n} \right). \tag{A1}$$

The matrix inside parentheses is the *centring matrix* (projecting onto the complement of $\mathbf{1}$), with eigenvalues 1 (multiplicity $n-1$) and 0 (eigenvector $\mathbf{1}$). The spectrum of $\mathbb{E}[S_A]$ is therefore $\{-n\mu_A$ (mult. $n-1$), 0 (eigenvector $\mathbf{1}$)$\}$.

#### HOI sector: $\mathbb{E}[\boldsymbol{S_{\hat{B}}}]$

Recall that $\tilde{B}_{ijk} \sim \mathcal{N}(\mu_B, n^{-2})$, with the zero-slice-sum constraint $\tilde{B}_{iii} = -\sum_{(j,k)\neq(i,i)} \tilde{B}_{ijk}$.

***Off-diagonal* $(i \neq j)$*.***
When $i \neq j$, none of the four summands in $(S_{\hat{B}})_{ij}$ hits a triple-diagonal entry. All entries are therefore unconstrained draws of $\tilde{B}$, each of mean $\mu_B$; with $4n$ terms in total and a $\frac{1}{4}$ prefactor,

$$\mathbb{E}[(S_{\hat{B}})_{ij}] = \tfrac{1}{4}(4n\mu_B) = n\mu_B.$$

***Diagonal* $(i = j)$*.***
Collecting identical pairs: $(S_{\hat{B}})_{ii} = \frac{1}{2}\sum_k (\tilde{B}_{iik} + \tilde{B}_{iki})$. Consider $\sum_k \tilde{B}_{iik}$; splitting at $k = i$:

$$\sum_k \tilde{B}_{iik} = \tilde{B}_{iii} + \sum_{k\neq i} \tilde{B}_{iik}.$$

The first term is the negative of $n^2-1$ draws of mean $\mu_B$; the second is $n-1$ draws. Their expectations sum to $-(n^2-1)\mu_B + (n-1)\mu_B = -n(n-1)\mu_B$. By an identical argument with indices two and three swapped, $\mathbb{E}[\sum_k \tilde{B}_{iki}] = -n(n-1)\mu_B$. Therefore

$$\mathbb{E}[(S_{\hat{B}})_{ii}] = \tfrac{1}{2}\big(-n(n-1)\mu_B - n(n-1)\mu_B\big) = -n(n-1)\mu_B.$$

Combining:

$$\mathbb{E}[S_{\hat{B}}] = n\mu_B\,\mathbf{1}\mathbf{1}^\top - n^2\mu_B\,I,$$

which factors as

$$\mathbb{E}[S_{\hat{B}}] = -n^2\mu_B\left(I - \frac{\mathbf{1}\mathbf{1}^\top}{n}\right). \tag{A2}$$

The spectrum of $\mathbb{E}[S_{\hat{B}}]$ is $\{-n^2\mu_B \text{ (mult. } n{-}1),\ 0 \text{ (eigenvector } \mathbf{1})\}$.

##### Parallel structure

Equations (A1) and (A2) share the same centring-matrix form. The two sectors differ only in the scalar prefactor ($-n\mu_A$ versus $-n^2\mu_B$), reflecting the different combinatorial footprint of pairwise versus three-way interactions: a pairwise row aggregates $O(n)$ entries whereas an HOI slice aggregates $O(n^2)$. In both cases, facilitative means ($\mu_A, \mu_B > 0$) produce a negative spike in the $(n{-}1)$-dimensional subspace orthogonal to $\mathbf{1}$ (stabilising), while competitive means ($\mu_A, \mu_B < 0$) produce a positive spike (destabilising). This is the structural basis for the regime-dependent choice of spectral thresholds in Sec. 4.

#### Appendix B Scaling of the spectral thresholds

Both $\delta_A$ and $\delta_B$ are determined by top eigenvalues of random symmetric matrices with zero-sum diagonal structure; the only differences are the entry variance and, when $\mu_A \neq 0$ or $\mu_B \neq 0$, the deterministic mean-field spike derived in Appendix A. We derive the common scaling via Weyl's inequality, the semicircle law, and the asymptotics of Gaussian maxima, treating both sectors in parallel.

***General strategy.***

For $S \in \{S_A, S_{\hat{B}}\}$, we decompose $S = \mathbb{E}[S] + \tilde{S}$, where $\tilde{S}$ is the centred (zero-mean) fluctuation. In the baseline and facilitative regimes the threshold is set by $\lambda_{\max}(\tilde{S})$; in the competitive regime it is set by $\lambda_{\max}(S)$ itself, which is dominated by the mean-field spike. For the centred part we further decompose $\tilde{S} = W + D$ into a hollow (zero-diagonal) Wigner matrix $W$ and a diagonal $D$ inherited from the zero-sum shift. Weyl's inequality gives the upper bound $\lambda_{\max}(\tilde{S}) \le \lambda_{\max}(W) + \lambda_{\max}(D)$, and we bound each term separately using the semicircle law and the Gaussian-maximum asymptotic [23]:

$$\max_{1\le i\le n} |Z_i| = \sigma_D\sqrt{2\log n}\,(1+o(1)) \qquad \text{a.s. as } n\to\infty, \tag{B3}$$

for i.i.d. $Z_i \sim \mathcal{N}(0, \sigma_D^2)$.

##### Pairwise threshold $\boldsymbol{\delta_A}$

***Baseline and facilitative regimes ($\mu_A \ge 0$).***

We compute $\lambda_{\max}(\tilde{S}_A)$.

*Off-diagonal (Wigner) contribution.* For $i \neq j$, $(\tilde{S}_A)_{ij} = \frac{1}{2}(\tilde{A}_{ij} + \tilde{A}_{ji}) - \mu_A$. Since $\tilde{A}_{ij}$ and $\tilde{A}_{ji}$ are independent draws of variance $1/n$, $\mathrm{Var}((\tilde{S}_A)_{ij}) = 1/(2n)$. The hollow part $W_A$ is therefore a Wigner matrix with upper-triangular variance $1/(2n)$, and the semicircle law gives $\lambda_{\max}(W_A) \xrightarrow{a.s.} \sqrt{2}$.

*Diagonal contribution.* The diagonal of $\tilde{S}_A$ is $\tilde{A}_{ii} + (n-1)\mu_A$. Since $\tilde{A}_{ii} = -\sum_{j\neq i} \tilde{A}_{ij}$ is a sum of $n-1$ independent $\mathcal{N}(\mu_A, n^{-1})$ draws, the centred diagonal has variance $(n-1)/n \to 1$. Applying (B3) with $\sigma_D = 1$ gives $\lambda_{\max}(D_A) = \sqrt{2\log n}\,(1+o(1))$.

Combining via Weyl's inequality:

$$\delta_A = \lambda_{\max}(\tilde{S}_A) \leq \sqrt{2} + \sqrt{2\log n}\,(1+o(1)) = O(\sqrt{\log n}). \tag{B4}$$

***Competitive regime ($\mu_A < 0$).***

The $(n-1)$-fold eigenvalue of $\mathbb{E}[S_A]$ is $+n|\mu_A| > 0$, producing a destabilising spike that dominates $\lambda_{\max}(S_A)$. In this regime we set

$$\delta_A = \lambda_{\max}(S_A) \sim n|\mu_A| = O(n), \tag{B5}$$

so that the threshold absorbs the mean-field spike.

#### HOI threshold $\delta_B$

***Baseline and facilitative regimes ($\mu_B \geq 0$).***

We compute $\lambda_{\max}(\tilde{S}_{\hat{B}})$.

*Off-diagonal (Wigner) contribution.* For $i \neq j$, $(\tilde{S}_{\hat{B}})_{ij} = (S_{\hat{B}})_{ij} - n\mu_B$ is $\frac{1}{4}$ times a sum of $4n$ independent centred draws each of variance $n^{-2}$, so $\mathrm{Var}((\tilde{S}_{\hat{B}})_{ij}) = \frac{1}{16} \cdot 4n \cdot n^{-2} = 1/(4n)$. The hollow part $W_B$ is a Wigner matrix with upper-triangular variance $1/(4n)$, and the semicircle law gives $\lambda_{\max}(W_B) \xrightarrow{a.s.} 1$.

*Diagonal contribution.* From Appendix A, $(S_{\hat{B}})_{ii}$ involves $2n(n-1)$ independent $\mathcal{N}(\mu_B, n^{-2})$ draws with a $\frac{1}{2}$ prefactor, so the centred diagonal has variance $\frac{1}{4} \cdot 2n(n-1) \cdot n^{-2} = (n-1)/(2n) \to 1/2$. Applying (B3) with $\sigma_D = 1/\sqrt{2}$ gives $\lambda_{\max}(D_B) = \sqrt{\log n}\,(1+o(1))$.

Combining via Weyl's inequality:

$$\delta_B = \lambda_{\max}(\tilde{S}_{\hat{B}}) \leq 1 + \sqrt{\log n}\,(1+o(1)) = O(\sqrt{\log n}). \tag{B6}$$

***Competitive regime ($\mu_B < 0$).***

The $(n-1)$-fold eigenvalue of $\mathbb{E}[S_{\hat{B}}]$ is $+n^2|\mu_B| > 0$, dominating $\lambda_{\max}(S_{\hat{B}})$. We set

$$\delta_B = \lambda_{\max}(S_{\hat{B}}) \sim n^2|\mu_B| = O(n^2). \tag{B7}$$

#### Numerical verification

Figure B1 verifies the bounds numerically across all regimes. In the baseline and facilitative regimes, both thresholds exhibit the predicted $O(\sqrt{\log n})$ scaling and the Weyl bound holds exactly for every replicate. In the competitive regime, the mean-field spike produces the expected $O(n)$ scaling for $\delta_A$ and $O(n^2)$ scaling for $\delta_B$. Ecologically, the $n$-power gap between sectors reflects a basic constraint: predominantly competitive communities require self-regulation to prevent competitive exclusion [6, 7].

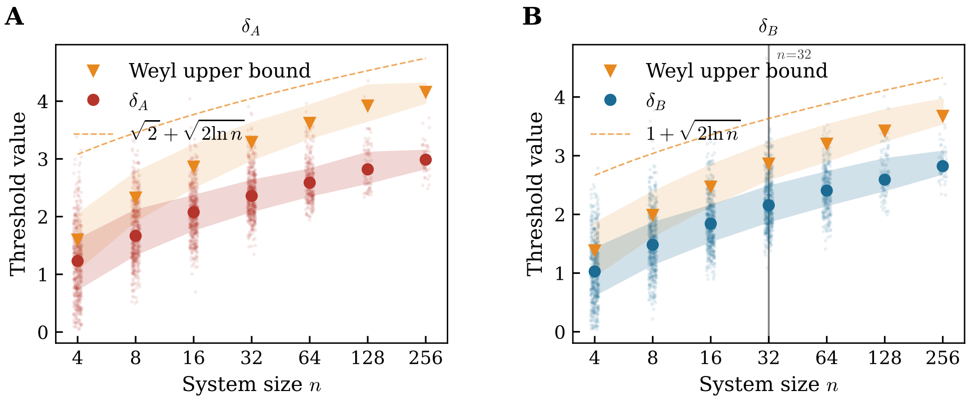


**Fig. B1** Numerical verification of the Weyl upper bound on the spectral thresholds $\delta_A$ and $\delta_B$ across the three regimes. Each matrix is decomposed into its hollow Wigner part $W$ and the diagonal $D$ inherited from the zero-sum shift (baseline and facilitative regimes), or evaluated directly (competitive regime). Weyl's inequality gives the exact (non-asymptotic) bound $\lambda_{\max}(\tilde{S}) \le \lambda_{\max}(W) + \lambda_{\max}(D)$ in the first two regimes, shown by orange triangles (medians) and shaded bands (inter-quartile ranges). Semi-transparent dots are individual replicates; filled circles and coloured bands show their medians and IQRs. The dashed lines are the asymptotic upper bounds from Eqs. (B4)–(B7). The exact Weyl bound holds for every replicate at every $n$ in the baseline and facilitative regimes; in the competitive regime, the $O(n)$ and $O(n^2)$ scalings of $\delta_A$ and $\delta_B$ are recovered.

### Appendix C Derivation of $\alpha_{eff}^{Taylor}$

Differentiating $f_i$ once yields

$$J_{ij} \;=\; (1-\alpha)\,A_{ij} \;+\; \alpha \sum_{l=1}^{n} \big(B_{ijl} + B_{ilj}\big). \tag{C8}$$

Summing over $j$ with $x_j^\star = 1$ and applying Eq. (**??**),

$$\begin{aligned} T_1[i] \;&=\; \sum_j J_{ij} \;=\; (1-\alpha)\sum_j A_{ij} \;+\; \alpha \sum_{j,l}\big(B_{ijl}+B_{ilj}\big) \\ &=\; (1-\alpha)(-1) \;+\; 2\alpha\,(-1) \;=\; -(1+\alpha). \end{aligned} \tag{C9}$$

Differentiating a second time gives

$$H_{ijk} \;=\; \alpha\,\big(B_{ijk} + B_{ikj}\big), \tag{C10}$$

so that at $\mathbf{x}^\star = \mathbf{1}_n$,

$$T_2[i] \;=\; \tfrac{1}{2}\sum_{j,k} H_{ijk} \;=\; \tfrac{\alpha}{2}\sum_{j,k}\big(B_{ijk}+B_{ikj}\big) \;=\; \alpha \sum_{j,k} B_{ijk} \;=\; -\alpha, \tag{C11}$$

where the penultimate equality uses the relabeling $j \leftrightarrow k$, and the last equality uses the slice-sum constraint in Eq. (**??**). Both $T_1[i]$ and $T_2[i]$ are independent of $i$, so averaging their absolute values over species gives, for $\alpha \in [0,1]$,

$$P \;=\; 1+\alpha, \qquad Q \;=\; \alpha, \tag{C12}$$

and substituting into Eq. (35) yields Eq. (36).

### Appendix D Regimes in Gibbs et. al (2024)

Here we describe in detail the parameterizations used for the robustness check, which are the same used by Gibbs et al. All parameterizations use $S = 20$ species.

***Regime Q1 (Dirichlet-constrained HOIs, varying pairwise strength).***

Off-diagonal entries of $A$ are drawn from Normal$(\mu_A, \sigma_A)$ with self-regulation fixed at $A_{ii} = 1$. Two parameter sweeps are combined: (i) $\mu_A \in [-0.75, 0.25]$ with $\sigma_A \in \{0, 0.1\}$; (ii) $\mu_A \in \{-1/S, 0, 1/S\}$ with $\sigma_A \in [0.7/\sqrt{S}, 1.6/\sqrt{S}]$. The target equilibrium is $\mathbf{x}^\star = (R_1/A_{11})\,\mathbf{1}$ (unit carrying capacity when $R_i = A_{ii} = 1$). HOI coefficients $B_{ijk}$ are determined by distributing the per-species residual $R_i - \sum_j A_{ij} x_j^\star$ across distinct pairs $(j, k)$ with $j \neq i$, $k \neq i$ using symmetric Dirichlet weights, so that all $B_{ijk}$ share the sign of the residual. When pairwise interactions are net facilitative this yields non-negative (competitive) HOIs, corresponding to the high-stability region identified in Fig. 2 of Gibbs et al.

***Regime Q2 (varying target abundances).***

Two sub-grids are combined. (i) *Specified abundance*: $\mu_A \in \{-0.5/S, 0.5/S\}$, $\sigma_A = 0.15$, and the target equilibrium is a constant vector $\mathbf{x}^\star = x_0\,\mathbf{1}$ with $x_0 \in [0.01, 1.99, 10]$; HOI coefficients are sampled via the same Dirichlet construction as Q1. (ii) *Noisy carrying capacity*: $\mu_A = 0.5/S$, $\sigma_A \in \{0, 1/\sqrt{S}\}$, and $\mathbf{x}^\star$ is drawn near the carrying capacity $R_i/A_{ii}$ with additive Gaussian noise of standard deviation $\eta \in$ linspace(0, 0.75, 10), clipped to a minimum of 0.01 and rescaled to preserve the mean; HOI coefficients use a non-equal-abundance Dirichlet construction in which each weight is divided by $x_j^\star x_k^\star$. In both sub-grids, $A_{ii} = 1$ and HOI coefficients can take either sign, corresponding to the moderate-stability region in Fig. 3 of Gibbs et al.

***Regime Q3 (correlation-stabilized, strong interactions).***

Off-diagonal entries of $A$ are drawn with $\mu_A \in \{-0.5/S, 0, 0.5/S\}$ and $\sigma_A \in$ linspace(0.5, 1.5, 3), allowing strong pairwise interactions. The aggregated HOI effect matrix $\tilde{B}_{ij} = \sum_k (B_{ijk} + B_{ikj}) x_k^\star$ is constructed as

$$\tilde{B}_{ij} = -p\,A_{ij} + (1-p)\,Y_{ij}, \tag{D13}$$

where the random component $Y_{ij}$ is initialized from Uniform$(0, 1/S)$ and then adjusted to satisfy the row-sum constraint $\sum_j \tilde{B}_{ij} = 2(R_i - \sum_j A_{ij})$, with anti-correlation strength $p \in$ linspace(0.1, 1.0, 10). This ensures that the HOI and pairwise contributions to the Jacobian have opposing sign structure—a mechanism that confers near-certain stability even with strong interactions (Fig. 4c of Gibbs et al.). The full tensor $B_{ijk}$ is obtained by distributing $\tilde{B}_{ij}$ across the $k$-indices.

### Appendix E Full simulation results figures

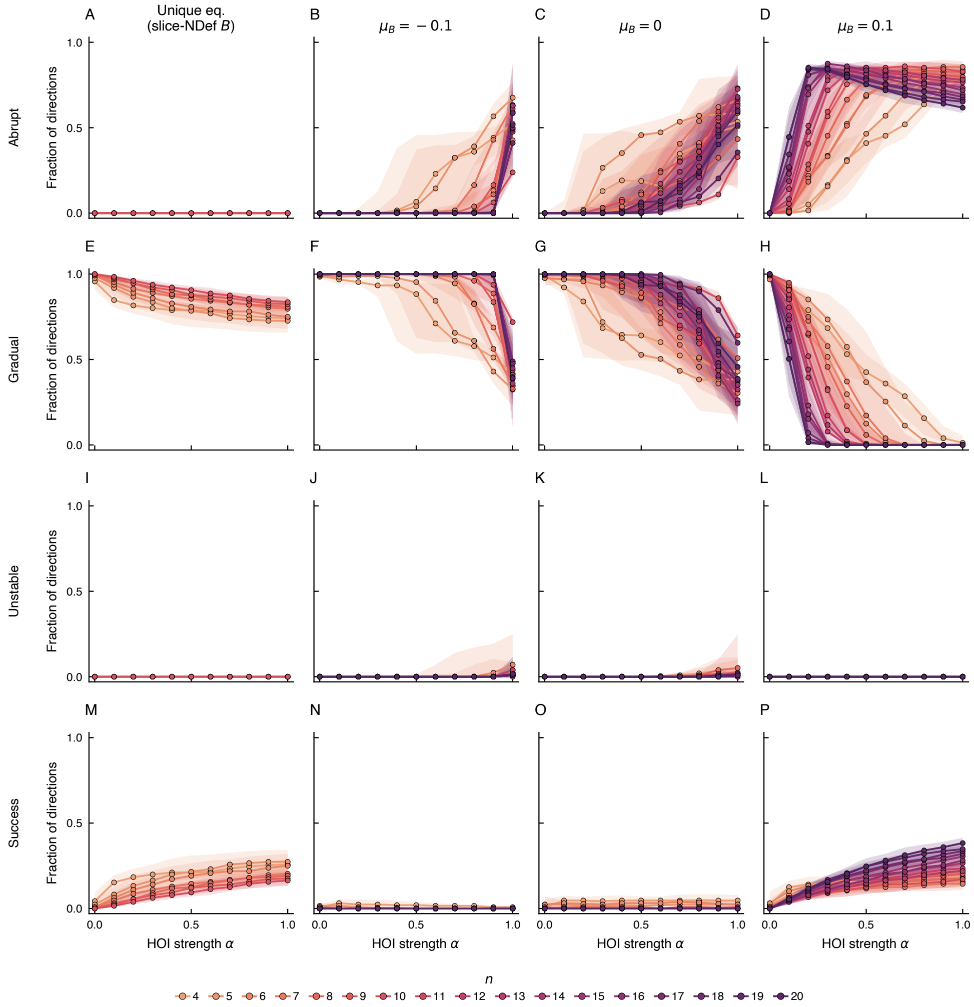


**Fig. E2** Fraction of perturbation directions leading to each boundary type as a function of HOI strength $\alpha$, across community sizes $n = 4$–$20$. Columns correspond to four simulation banks: unique equilibrium parameterization, $\mu_B = -0.1$, $\mu_B = 0$, and $\mu_B = 0.1$. Rows show the fraction of directions classified as abrupt (**A**–**D**), gradual (**E**–**H**), unstable (**I**–**L**), and successful (**M**–**P**). Lines indicate medians across model replicates; shaded bands span the interquartile range. Colours encode community size $n$. As $\alpha$ increases, the fraction of abrupt boundaries grows while gradual boundaries decline, with the transition sharpening for larger $n$ and positive $\mu_B$. The successful-direction fraction increases monotonically with $\alpha$ and $n$ for the unique equilibrium and facilitative parameterizations, while it remains very low for the competitive and mixed parameterizations.

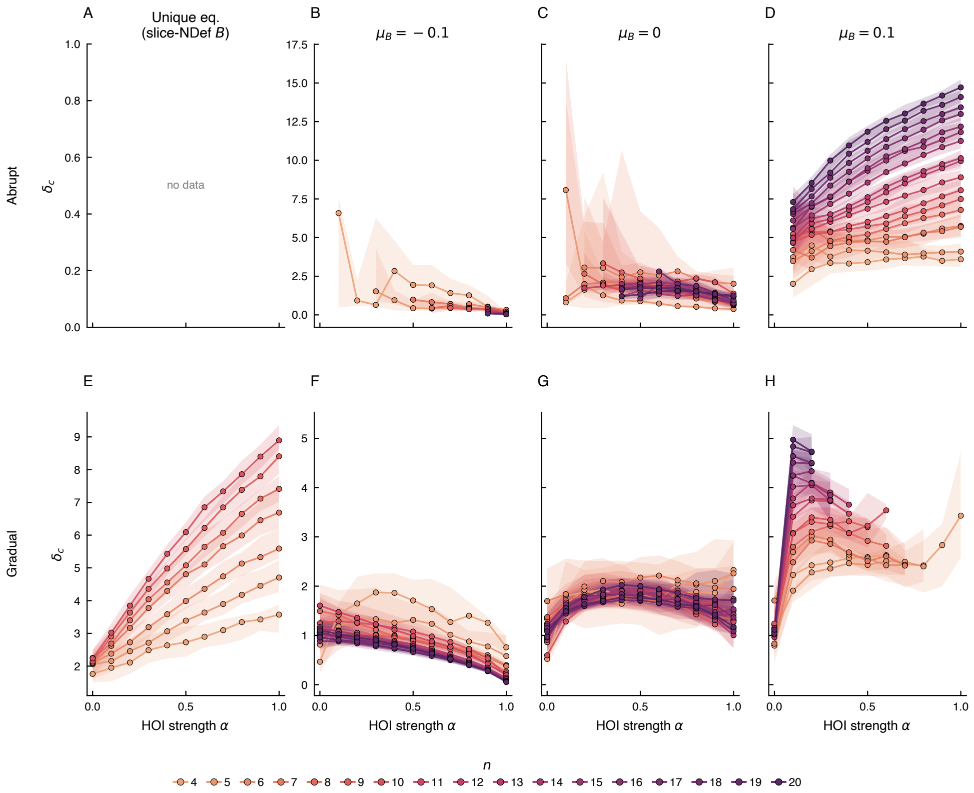


**Fig. E3** Critical perturbation magnitude $\delta_c$ as a function of HOI strength $\alpha$, across community sizes $n = 4$–$20$. Columns correspond to four simulation banks: unique equilibrium, $\mu_B = -0.1$, $\mu_B = 0$, and $\mu_B = 0.1$. Rows show $\delta_c$ for abrupt (**A**–**D**) and gradual (**E**–**H**) Lines indicate medians across model replicates; shaded bands span the interquartile range. Colours encode community size $n$. Panel **A** display "no data" because the abrupt boundary type is absent for the unique equilibrium parameterization.

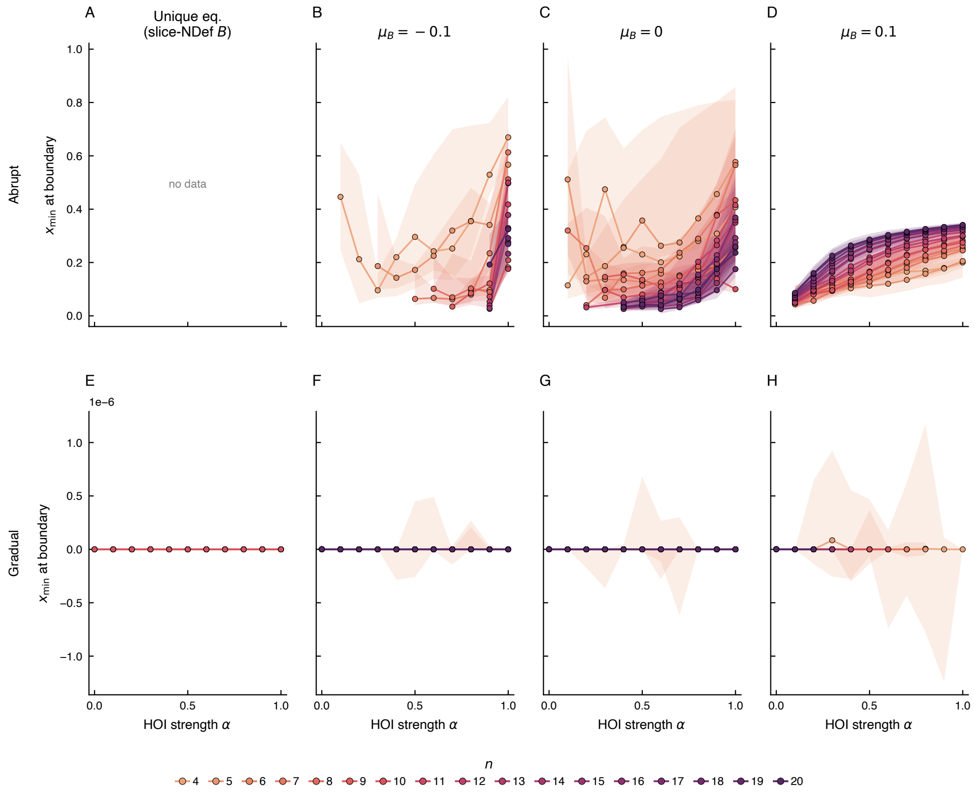


**Fig. E4** Minimum species abundance $x_{\min}$ at the tipping boundary as a function of HOI strength $\alpha$, across community sizes $n = 4$–20. Columns correspond to four simulation banks: unique equilibrium, $\mu_B = -0.1$, $\mu_B = 0$, and $\mu_B = 0.1$. Rows show $x_{\min}$ for abrupt (**A**–**D**) and gradual (**E**–**H**) boundary types. Lines indicate medians across model replicates; shaded bands span the interquartile range. Colours encode community size $n$. For abrupt boundaries, $x_{\min}$ remains strictly positive, confirming that species abundances have not yet reached zero when the fold bifurcation occurs. For gradual boundaries, $x_{\min}$ is effectively zero ($\sim 10^{-6}$), consistent with the boundary being defined by a species abundance crossing zero.

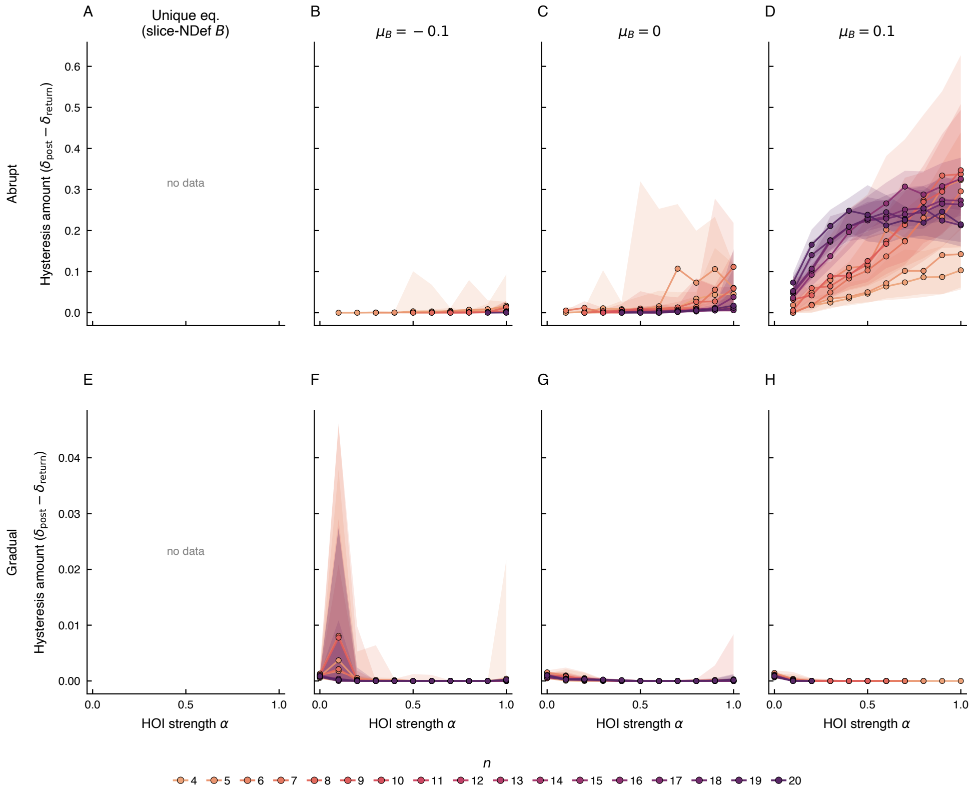


**Fig. E5** Hysteresis amount ($\delta_c - \delta_{\mathrm{recover}}$) as a function of HOI strength $\alpha$, across community sizes $n = 4$–$20$. Columns correspond to four simulation banks: unique equilibrium, $\mu_B = -0.1$, $\mu_B = 0$, and $\mu_B = 0.1$. Rows show hysteresis for abrupt (**A**–**D**) and gradual (**E**–**H**) boundary types. The hysteresis amount measures the difference between the forward critical perturbation and the perturbation at which the system returns to its original state upon reversal. Lines indicate medians across model replicates; shaded bands span the interquartile range. Colours encode community size $n$. Abrupt boundaries exhibit hysteresis while gradual ones do not, consistent with the irreversibility expected from fold bifurcations.
